# StabCell: Stability selection for clustering and marker detection in single-cell RNA sequencing

**DOI:** 10.64898/2026.05.07.720061

**Authors:** Niklas Lück, Andrea Rossi, Christian Staerk

## Abstract

Conventional pipelines for differential expression analysis in single-cell RNA sequencing (scRNA-seq) data first cluster individual cells and then test for differentially expressed genes between the resulting clusters. Using the same data for clustering and testing, however, poses a selective inference problem and can result in overconfidence in differences that may not reflect true biological variation. We introduce *StabCell*, a stability selection framework which integrates clustering and detection of differentially expressed marker genes. By repeatedly performing clustering and differential expression analysis on complementary pairs of random subsamples, *StabCell* assesses clustering and marker stability, yielding a stable clustering, cell-specific stability scores, and sets of stable marker genes. In simulations, we demonstrate that *StabCell* generally maintains empirical per-family error rate (PFER) below prespecified thresholds, selecting fewer false positive marker genes than the conventional approach, especially in cases with low signal-to-noise ratio and low sequencing depth. Simultaneously, *StabCell* provides competitive marker detection performance compared to alternative approaches. An application to a cell differentiation dataset from induced pluripotent stem cells (IPSCs) to cardiomyocytes illustrates that *StabCell* can reduce overclustering and prioritize biologically meaningful marker genes, improving the robustness and interpretability of scRNA-seq analyses.

## 1 Introduction

Single-cell RNA sequencing (scRNA-seq) has given researchers unparalleled insight into biological tissues and their cellular mechanisms and is widely used to investigate diverse research questions (Zappia and Theis, 2021). A common goal is to determine the cell-type composition of the tissue under study. Since manual annotation of cells quickly becomes infeasible as datasets grow in size, scalable computational approaches are needed. Typically, cells are first clustered to identify cellular subpopulations (Kiselev et al., 2019; Yang et al., 2019; Zhang et al., 2024). The resulting clusters are then compared using statistical tests to identify genes that are differentially expressed between subpopulations, so-called marker genes, which can be used to annotate clusters (Li et al., 2022; Zhang et al., 2026).

Conventionally, such analyses use the same data for clustering and for differential expression testing, which can result in overconfident *p*-values. This selective inference problem, also known as “double dipping”, has been highlighted among the eleven grand challenges in single-cell data science (Lähnemann et al., 2020). Although this problem has been recognized for some time, popular software packages like Seurat (Hao et al., 2024) and scanpy (Wolf et al., 2018) still use this approach, with warnings such as “*p*-values should be interpreted cautiously” (Satija et al., 2026).

In related situations, statisticians randomly split the observations into different subsets for each task. Here, however, a simple split of cells into a clustering and a marker detection set does not solve the double dipping problem, since cluster labels are also required for marker detection and a naive label transfer (e.g. by kNN classification) from one subset to the other also transfers information (Chen and Witten, 2023). TN Test (Zhang et al., 2019) is an approach which manages to incorporate observation splitting by fitting a hyperplane separating two clusters in the clustering data, using it to assign cluster labels in the marker data, and performing a truncated normal test. Count splitting is a form of data thinning that can avoid this information leakage under suitable assumptions (Neufeld et al., 2024a,b) by splitting the matrix of observed count values into two independent matrices of the same dimensions. Taking the sum of both matrices recovers the original data.

Overconfident *p*-values can arise from spurious subpopulations introduced by the clustering method. Such over-clustering may lead to spurious marker gene selection and misleading biological interpretations. Methods such as recall (DenAdel et al., 2025) use artificial variables to reduce overclustering. Similarly, ClusterDE (Song et al., 2026) uses synthetic null data to control the false discovery rate (FDR) of selected marker genes when comparing two clusters.

Other approaches like singleCellHaystack (Vandenbon and Diez, 2020), scMEB (Zhu and Yang, 2023), SEMI-TONES (Vlot et al., 2022), MarcoPolo (Kim et al., 2022), and scTIM (Feng et al., 2020) can detect sets of marker genes without prior cluster label assignment. The latter method is subsample-based and identifies markers by consensus optimization. These markers are then used in a second step for hierarchical clustering to group similar cells into subpopulations.

In this article, we introduce *StabCell*, a framework that jointly improves the robustness of clustering and marker gene selection while using information from all cells. *StabCell*’s main idea is to borrow concepts from stability selection (Meinshausen and Bühlmann, 2010; Shah and Samworth, 2013), a framework that can provide error control for variable selection under suitable assumptions, and apply them to marker gene selection across independently clustered subsamples of the full data. Our algorithm generates random pairs of complementary subsamples, where each subsample of cells is independently clustered and analyzed for differential expression. Aggregation of all subsample clusterings provides a stable clustering of the full dataset and cell-specific stability scores. Similarly, the selected marker genes in the subsamples are used to obtain cluster-specific sets of stable markers. We compared *StabCell* against the conventional double dipping pipeline and competing methods in simulations and on experimental data from induced pluripotent stem cells (IPSCs) differentiating into cardiomyocytes. Simulation results show that *StabCell* generally maintains empirical per-family error rate (PFER) below prespecified thresholds and quantifies the stability of cluster assignments for individual cells. The real-data application demonstrates its ability to reduce overclustering and to prioritize biologically meaningful genes.

## 2 Methods

*StabCell* is designed to infer a stable clustering of scRNA-seq data and to identify stable marker genes for these clusters (Fig. 1). Complementary subsamples of the cells are generated and independently analyzed. Aggregating the results across subsamples yields a stable clustering, cell-specific stability scores, and sets of stable marker genes. Here we focus on the widely used Seurat (Hao et al., 2024) analysis pipeline for clustering and differential expression; however, our approach is modular and can, in principle, be combined with any alternative clustering and marker detection method. Full algorithmic details are provided in Supplementary Algorithm S1.

**Figure 1.**
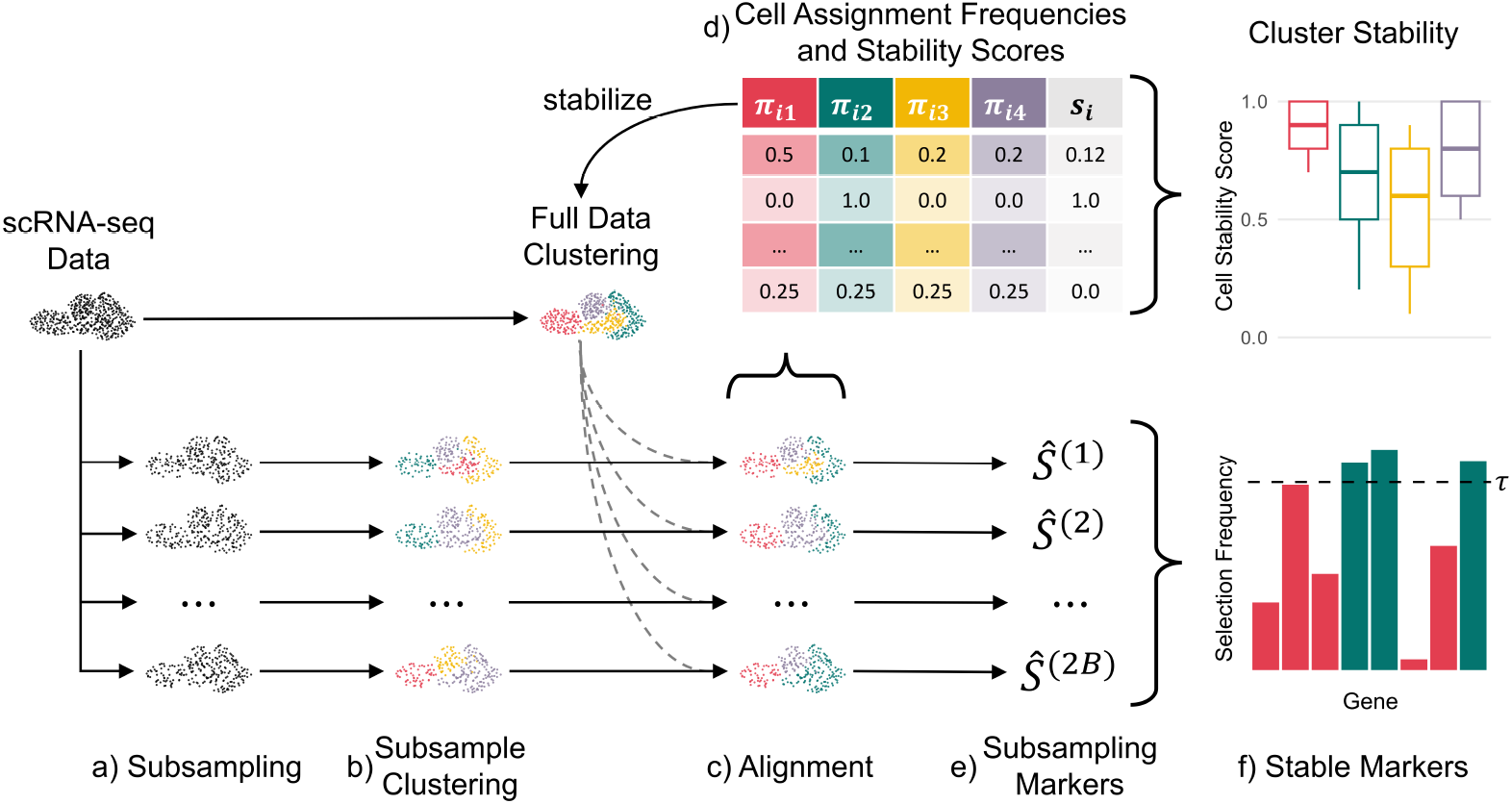
Overview of *StabCell*: a) The dataset is split *B* times into pairs of complementary subsamples. b) For each subsample, independent clustering is performed. c) Subsample clusterings *η*^(*b*)^ (*b* = 1, … , 2*B*) are aligned to an initial clustering *η*^(0)^ of the full data. d) Cell assignment frequencies *π*_*ik*_ to each cluster *k* = 1, … , *K*_0_ are calculated. They are used to calculate cell stability scores *s*_*i*_ and to infer a stable clustering *η*^(stable)^ of the full dataset. e) After an additional alignment step, each subsample cluster is investigated for marker genes *Ŝ*^(*b*)^. f) Genes with selection frequencies at least *τ* across the subsamples are selected as stable marker genes.

### 2.1 Stable Clustering

*StabCell* starts by generating *B* complementary pairs of non-overlapping subsamples *D*^(*b*)^ from the full scRNA-seq dataset *D* with *n* cells and *d* genes. Each subsample *D*^(*b*)^ contains 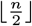 cells, yielding a total of 2*B* subsamples. Let *I*^(*b*)^ *⊆ {*1, … , *n}* denote the index set of cells contained in subsample *D*^(*b*)^ for *b* = 1, … , 2*B*.

A clustering approach *f*_*c*_ is used to obtain clusterings *η*^(*b*)^ = *f*_*c*_ *D*^(*b*)^ for each subsample, where a clustering vector *η*^(*b*)^ denotes the cluster labels 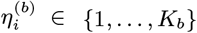 for *i ∈ I*^(*b*)^. Potential clustering methods include the commonly used Louvain (Blondel et al., 2008) or Leiden (Traag et al., 2019) algorithms for community detection in the cells’ *k*-nearest neighbor graph. An initial clustering 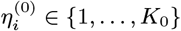 of the full dataset (which in the simples case is generated as *η*^g(0)^ = *f*_*c*_ (*D*)) serves as a reference to align the subsample clusterings.

Every cluster *k* in each subsample *b* is aligned to its closest cluster *k*^*′*^ in *η*^(0)^. Considering a cluster *k* in clustering *η*^(*b*)^, we denote its set of cell indices by 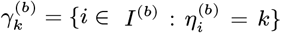. Analogously, 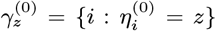 denotes the set of cells assigned to cluster *z* in the reference clustering. The similarity between two clusters in the different clusterings *η*^(*b*)^ and *η*^(0)^ is then measured by the Jaccard similarity of the index sets. Specifically, for cluster *k* in subsample clustering *η*^(*b*)^, the closest reference cluster *k*^*′*(*b*)^ is determined as:

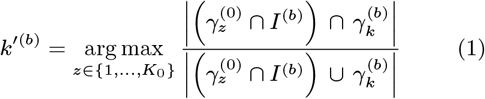

That is, each cluster *k* in subsample clustering *η*^(*b*)^ is mapped to the reference cluster *k*^*′*(*b*)^ in *η*^(0)^ with which it has the largest Jaccard similarity. Mapping cluster labels *k→ k*^*′*(*b*)^ yields the aligned clustering *η*^*′*(*b*)^. Note that each observation in each subset *D*^(*b*)^ gets assigned a cluster label from *η*^(0)^. Multiple clusters in *η*^(*b*)^ may be assigned the same cluster label from *η*^(0)^. Thus it is possible that clustering *η*^*′*(*b*)^ is composed of fewer clusters than *η*^(*b*)^ or *η*^(0)^ (see Supplementary Fig. S1 for an example).

All aligned cluster labels 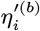 of cell *i* are used to calculate its assignment frequency for reference cluster *k*^*′*^:

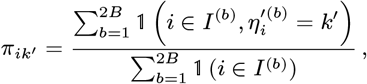

where 1 denotes the indicator function, which returns 1 if the specified conditions are true and 0 otherwise. Whenever an even number of cells is analyzed, the denominator equals *B*.

A stable clustering *η*^(stable)^ is derived by assigning each observation to the cluster with maximal assignment frequency:

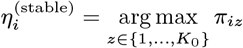

This can result in the merging of non-robust clusters from the reference clustering *η*^(0)^, which would decrease the overall number of clusters. In case of ties between two or more clusters with the same assignment frequency, a deterministic rule to pick one of the competing cluster labels is used. We break ties by selecting the cluster with the lowest numeric label. Note that the procedure cannot introduce new cluster labels beyond those present in *η*^(0)^. Therefore, an upper bound on the number of clusters in *η*^(stable)^ is determined by *η*^(0)^.

The assignment frequencies 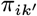 for all *K*_0_ *>* 1 clusters in *η*^(0)^ can also be used to obtain a cell stability score *s*_*i*_ for cell *i*:

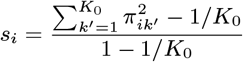

This score is based on the complement of the Gini impurity and is scaled to lie in the interval [0, 1]. If cell *i* is consistently assigned to the same cluster *k*^*′*^ in all subsamples in which it appears, i.e., if any 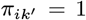, the score is 1, whereas it is 0 if the assignment frequencies are equal for all clusters (i.e., all 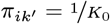).

### 2.2 Stable Marker Genes

Conditioning on the stable clustering *η*^(stable)^, we describe how to identify stable marker genes that differentiate one stable cluster from the rest of the cells.

A second Jaccard index-based alignment is performed for each cluster *l* in *η*^(stable)^ to identify the closest cluster *l*^*′*(*b*)^ in each subsample:

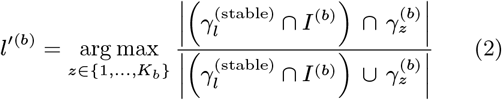

This step identifies a representative for reference cluster *l* in each subsample *b*, allowing corresponding marker genes to be detected across all 2*B* subsamples. Compared to Eq. (1), the arg max-operation now operates over the subsample clusterings *η*^(*b*)^ while keeping the cluster in the full data clustering *η*^(stable)^ constant. This can result in multiple clusters in *η*^(stable)^ being aligned to the same cluster in *η*^(*b*)^, or some clusters in *η*^(*b*)^ not being aligned to any cluster in *η*^(stable)^. Supplementary Fig. S1 highlights the differences between both cluster alignment steps.

The aligned cluster *l*^*′*(*b*)^ is tested against the union of all other clusters in *η*^(*b*)^ using a testing function *f*_*t*_ on the subset *D*^(*b*)^. A common choice to test for such differentially expressed genes is, e.g., the Wilcoxon rank-sum test. Bonferroni-adjusted *p*-values 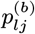 for cluster *l* and genes *j* = 1, … , *d* are used to construct the set of subsample markers 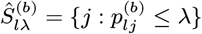. The *p*-value threshold *λ∈* (0, 1) controls the number of selected genes in the subsamples. We follow the testing convention and set *λ* = 0.05 if not specified differently. This threshold must not be interpreted as a statistical significance measure. It is purely a means to select candidate marker genes within the *StabCell* framework. Any marker selection procedure *f*_*t*_ can be used with *StabCell*. If the marker selection procedure returns a binary selection indicator, the resulting set of selected genes can be used without calculating *p*-values.

Selection frequencies 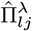 for each gene *j* are calculated utilizing all 2*B* sets of subsample markers:

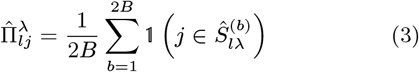

Finally, the set of stable marker genes 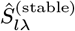^(stable)^ for cluster *l* is determined by thresholding the selection frequencies:

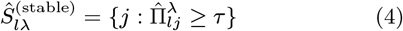

Stability selection (Meinshausen and Bühlmann, 2010) provides PFER control under exchangeability and additional assumptions on the base selection procedure. Shah and Samworth (2013) extended this framework using complementary subsamples and derived bounds for the expected number of selected variables with low selection probabilities under alternative assumptions (“none”, “unimodal”, and “*r*-concavity”). The bounds of Shah and Samworth (2013) concern genes with low selection probabilities, which do not necessarily coincide with truly non-differentially expressed genes. By specifying a nominal PFER_max_ target, we obtain the corresponding selection frequency threshold *τ* (Supplementary Section A).

We note that the results of Meinshausen and Bühlmann (2010) and Shah and Samworth (2013) assume that the base selection procedure is applied to the respective subsamples without using information from observations outside these subsamples. Since information from all cells is used for cluster alignment and generation of stable clustering *η*^(stable)^, information from cells not included in a subsample *D*^(*b*)^ could leak between subsamples. We therefore also propose an alternative variant of *StabCell* that mitigates this issue by splitting the cells into two parts (Supplementary Section B). Since it was shown to be more conservative (Supplementary Fig. S2) and the specified PFER_max_ thresholds were approximately maintained in most simulation settings (Section 3.1), we followed the original *StabCell* algorithm. However, the resulting error control should be interpreted as an empirically supported approximation and not as a direct finite-sample guarantee.

The presented method is designed to identify stable marker genes by comparing one cluster with the rest of the sample (1-vs-rest), but it can easily be adapted to detect stable markers 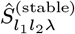 between a pair of clusters *l*_1_ and *l*_2_ (1-vs-1). This largely follows the same procedure with one modification. In the case that 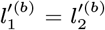, i.e., both clusters from the stable clustering are aligned with the same cluster of the subsample clustering, *p*-values 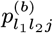 are set to 1 for all genes *j* since no difference between clusters 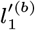 and 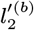 can be detected.

Furthermore, the PFER target is specified separately for each individual testing scenario. When, e.g., analyzing multiple clusters for stable 1-vs-rest markers, the over-all PFER target is the sum of the individually specified targets.

### 2.3 Computational Scalability

Applying the clustering and testing functions *f*_*c*_ and *f*_*t*_ to multiple subsamples increases computational cost. However, the main components of *StabCell* are embarrassingly parallel. In particular, the most computationally intensive operations are clustering and marker detection within the subsamples, which can be performed independently for each subsample. Only the aggregation of clustering and testing results is performed centrally. Thus, it is possible to reduce wall-clock time by leveraging parallel high-performance computing. Furthermore, the number of marker comparisons increases linearly with the number of clusters in *η*^(stable)^ for 1-vs-rest testing, while it increases quadratically for all pairwise 1-vs-1 comparisons. A runtime comparison between different methods is shown in Supplementary Fig. S3.

## 3 Results

We assessed *StabCell* in simulation studies and on a real scRNA-seq dataset of human cardiomyocyte differentiation. In all analyses, we used *B* = 50 complementary pairs of subsamples (i.e., 100 subsamples in total), as this provided a reasonable compromise between robustness and computational complexity. We considered all three bounds proposed by Shah and Samworth (2013) and we benchmarked *StabCell* against the double dipping approach, against Countsplit (Neufeld et al., 2024b; Poisson assumption with even split between clustering and testing), against ClusterDE (Song et al., 2026; Poisson assumption), and against TN Test (Zhang et al., 2019; 50 : 50 clustering-testing-split). These methods differ in their error control properties. For double dipping, Countsplit, and TN Test we used Bonferroni-adjusted *p*-values, with Countsplit and TN Test aiming to control the family-wise error rate (FWER). ClusterDE is designed to control the FDR. *StabCell*, however, aims to control the PFER, i.e., the expected number of false positives. For that reason, we focus on the number of true and false positive markers in our simulation study and also evaluate competing methods in that regard. For each of the competing methods we set the significance level to *α* = 5%.

Wherever it was permitted by an approach, we used functions from the `R` package `Seurat` (Hao et al., 2024) for clustering and testing with default parameters, unless stated otherwise. Since the simulated and experimental data were already available in a cleaned format, we did not perform additional cell or gene filtering for quality control in our clustering and testing functions, and instead considered the input count data matrix as fixed. The clustering procedure *f*_*c*_ consisted of data normalization, variable feature selection, data scaling, dimension reduction with PCA, kNN graph construction, and cluster identification using the Louvain algorithm (Blondel et al., 2008). For the testing function *f*_*t*_, we used a Wilcoxon rank-sum test without prefiltering of genes, yielding a *p*-value for each gene in each subsample.

There are different approaches to obtain the reference partition *η*^(0)^. The most straightforward is to apply the clustering function *f*_*c*_, intended for the subsamples, to the full dataset. Another approach we investigated was to use the co-clustering matrix across all subsample clusterings as a similarity matrix in hierarchical clustering. Comparing both approaches with the true partition in simulated settings revealed that the latter approach was, at best, comparable in terms of adjusted Rand index (ARI). Since it also requires a manual step to determine the height at which to cut the dendrogram, we prefer the first approach, where *η*^(0)^ = *f*_*c*_(*D*), and used it for all presented results.

### 3.1 Simulations

We used the `R` package `splatter` (Zappia et al., 2017) to simulate scRNA-seq datasets for two different scenarios: one where all observations were simulated from the same population, i.e., there were no true marker genes, and the other where multiple equally sized subpopulations were simulated. We varied the proportion of differentially expressed genes, i.e., the probability for each gene to be differentially expressed (“`de.prob`”, 1% and 5%), the location parameter controlling the strength of differential expression (“`de.facLoc`”, 0.1 and 0.4), and the library size (“`lib.loc`”, 10,000, 25,000, and 50,000 expected counts per cell before dropout). For each parameter setting, we simulated and analyzed 100 independent datasets with 5,000 cells and 10,000 genes to assess the robustness of the results.

To evaluate true marker recovery, we again used the Jaccard similarity to match each cluster in any approach to its most similar cluster in the simulated true cell clustering and used the respective true marker genes to calculate performance metrics. We then benchmarked *Stab-Cell* against the other approaches by comparing 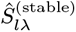 of cluster *l* in *η*^(stable)^ against the most similar one in the other approaches’ clusterings (cf. Supplementary Fig. S4).

#### 3.1.1 Two Cluster Simulation

In our first simulation setting, we simulated data from either one or two true cell populations and adaptively tuned the resolution parameter of the Louvain algorithm in our clustering function *f*_*c*_ to always obtain two clusters, thereby deliberately introducing a spurious cluster in the case of a single true cell population.

We investigated different settings for the nominal PFER target across all three assumptions. First, we focus on two true cell types with the most challenging scenarios of low probability and strength of differential expression (`de.prob` = 1%, `de.facLoc` = 0.1), where we observed that using either the unimodality-based threshold or the assumption-free threshold resulted in no false positive markers (Fig. 2). Under the *r*-concavity assumption, false positive markers are observed if the library size is low or if PFER_max_ is large, but even in those cases the observed mean is below the specified PFER threshold. The double dipping approach and Countsplit in comparison report more false positive marker genes. This reduction in false positives comes at the cost of reduced power to retrieve true positive marker genes and, as expected, the number of true positives is (depending on PFER_max_) on average lower when comparing *StabCell* to the double dipping approach. Other simulation settings with 2 true clusters show similar trends (Supplementary Figs. S5 and S6). ClusterDE and TN Test consistently retrieve few to no false positive markers. For true positive marker recovery, TN Test performs comparably to the two weaker assumptions at higher PFER_max_ values. However, in the same settings the *r*-concavity assumption retrieves more true positive markers. ClusterDE has two simulation settings in which its capability to retrieve true positives is similar to double dipping or Countsplit, but there are also some settings in which it performs markedly worse than all other approaches. Using different thresholds for *Stab-Cell*’s selection frequency or the *p*-value in other methods allowed us to compute the area under the precision recall curve (AUPRC) and under the receiver operating characteristic curve (AUROC). *StabCell* outperformed both Countsplit and TN Test in both metrics, while being comparable to double dipping (Supplementary Figs. S8 and S9).

**Figure 2.**
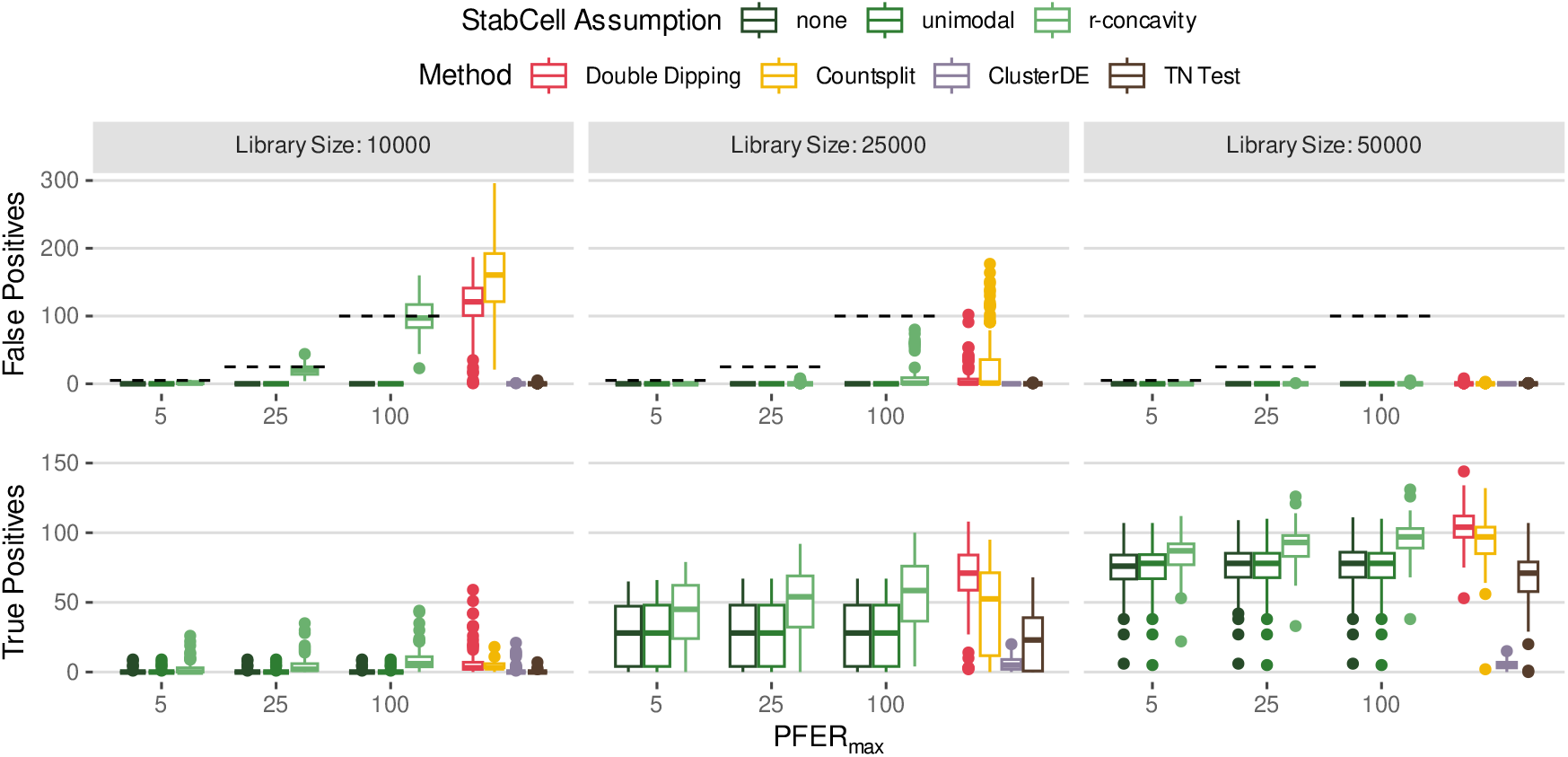
Marker detection results in terms of false (top) and true (bottom) positives for a differential expression probability of 1%, a differential expression strength of 0.1, and two true clusters. Boxplots are based on 100 independent simulation replicates. *StabCell* shows empirical PFER control. For other simulation settings, see Supplementary Figs. S5 to S7.

In the 1 cell type scenario, there are no true marker genes and every selected gene is a false positive. Under the unimodality and assumption-free bounds, once again no marker is selected across all PFER_max_ values (Supplementary Fig. S7). With increasing PFER_max_, the number of false positives grows under the *r*-concavity assumption. For low library sizes, the observed numbers of false positives cover the specified PFER_max_. For the lowest library size (10,000), the mean number of false positives is slightly larger than PFER_max_ thresholds 50 and 100. For larger library sizes, the number of observed false positives is consistently lower than PFER_max_. The *r*-concavity assumption generally yields the least conservative selection frequency threshold, and the observed mean number of false positives exceeded the specified PFER_max_ in only a few settings. Comparing *StabCell* to the other methods, *StabCell* selects fewer false positive markers than double dipping across the investigated PFER_max_ and for all assumptions. Countsplit only has fewer false positives for the combination of large library size (50,000) with large PFER_max_ (100) and *r*-concavity. ClusterDE and TN Test again have few to no false positive markers.

Beyond marker gene selection, *StabCell* can refine the reference clustering and provide cell-specific stability scores. In simulations with two true cell subpopulations, we observed improved ARI values compared to the other methods, provided that sufficient information was available (Fig. 3a for 1% differential expression probability and 0.1 differential expression strength). Similar trends were observed for *StabCell*’s mean cell stability scores, i.e., when little information was available to differentiate between clusters, or when only one true cell type was present, we observed lower cell stability scores (Fig. 3b). These findings are also consistent with the other simulation settings (Supplementary Figs. S10 and S11).

**Figure 3.**
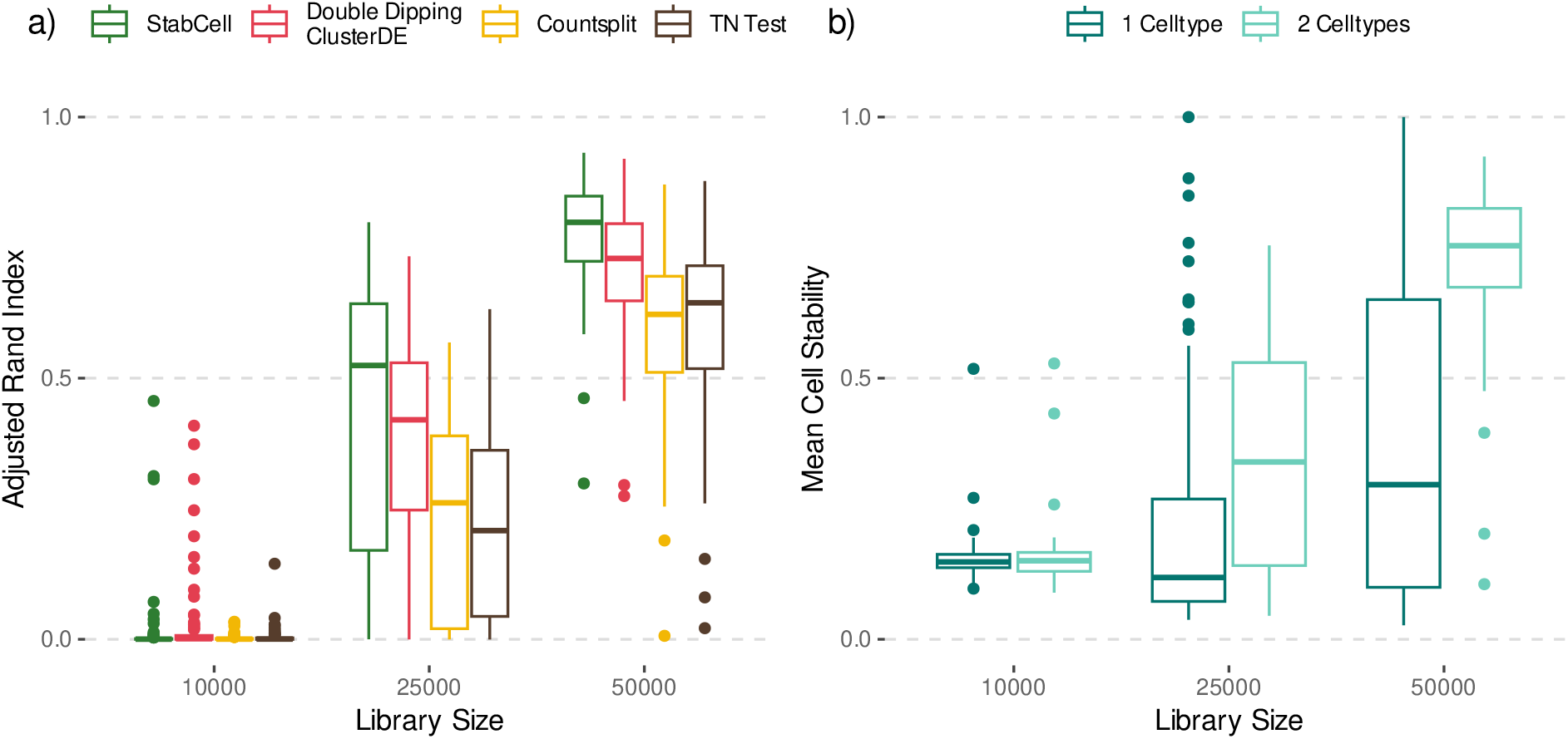
Clustering results for a differential expression probability of 1% and a differential expression strength of 0.1. Boxplots are based on 100 independent simulation replicates. a) Adjusted Rand Index (ARI) comparison between *StabCell* and other approaches in the two-cell-type setting. b) Comparison of mean cell stability scores between the one- and two-cell-type settings. For other differential expression settings, see Supplementary Figs. S10 and S11.

Investigating an individual example simulation setting with two moderately separated cell types reveals the informative value of the cell stability scores. Using UMAP (McInnes et al., 2020) for dimensionality reduction and coloring the cells by true simulated cell type results in overlapping clusters (Fig. 4). Coloring the cells by their stability score shows that especially cells in the boundary region are unstable regarding their cluster assignment. In contrast, the further away a cell is from the boundary, the more stable it becomes. When dividing the cells into groups with correct and incorrect cluster assignment, we observed that the initial clustering *η*^(0)^ had 773 incorrectly clustered cells, which *StabCell* reduced to 484 in *η*^(stable)^. We also see that the cells with wrong cluster assignment tended to have lower stability scores than correctly clustered cells.

**Figure 4.**
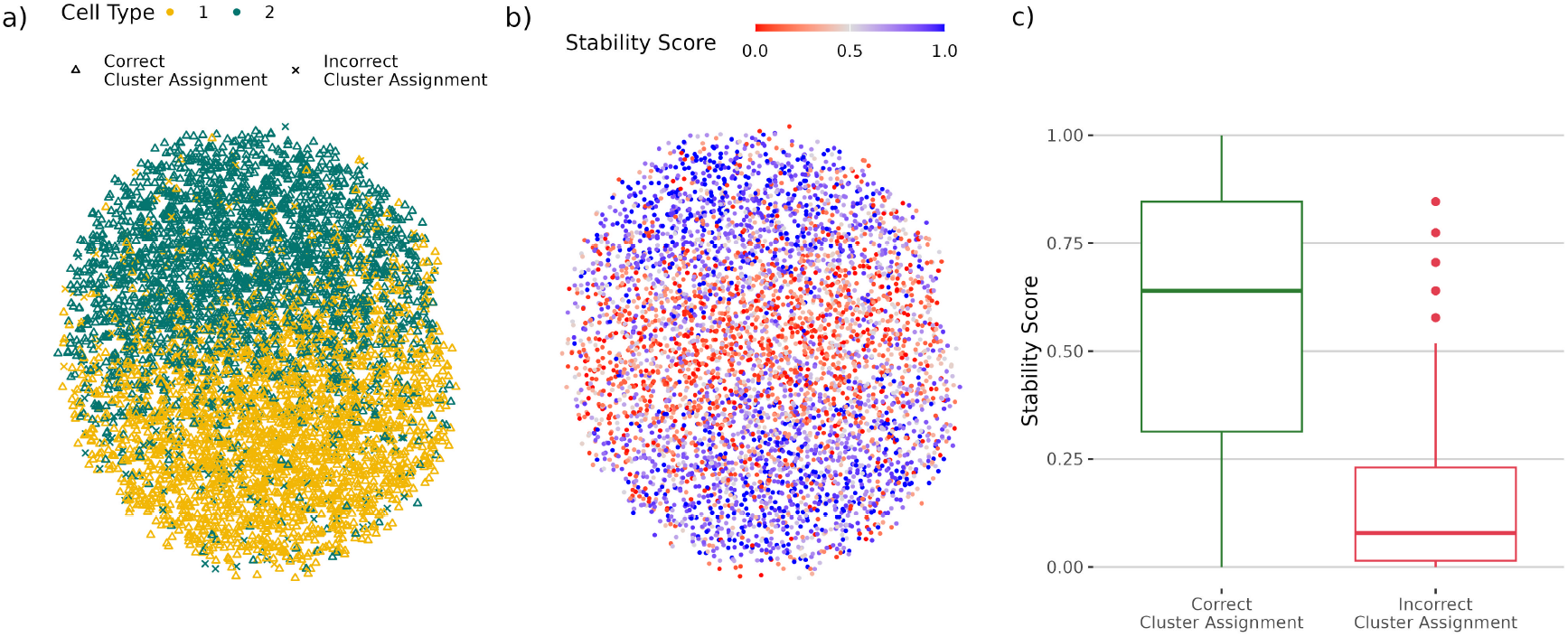
UMAP plots (McInnes et al., 2020) of an example simulation with two true cell types. a) Cells colored by their true simulated cell type with shapes corresponding to correct and incorrect cluster assignment in *η*^(stable)^. b) Cells colored by their stability score. c) Cells with incorrect cluster labels tend to have lower stability scores.

#### 3.1.2 Many Cluster Simulation

We performed another similar simulation study with either 1 or 10 true cell subpopulations and left the Louvain algorithm’s resolution parameter fixed at its default value of 0.8, allowing the number of detected clusters to vary across datasets and subsamples. This data-driven approach more closely resembles common bioinformatics practice. For all marker detection results, we selected the cluster with label “0” for ease of presentation.

Here, we compare one cluster against the union of all other clusters to determine differentially expressed genes. It should be noted, however, that methods ClusterDE and TN Test are designed to compare one cluster against another cluster. For comparability, we still use them with a 1-cluster-vs-rest test and therefore their results should be interpreted with caution.

False positive marker results are largely consistent with the specified PFER_max_. For the combination of 10 true cell types, 1% marker probability, a differential expression strength of 0.4, a large library size, and the *r*-concavity assumption, however, the mean number of false positives exceeded the specified PFER_max_ (Supplementary Fig. S13). While the *r*-concavity-based threshold generally performed well in many scenarios, it appeared too liberal in those few simulation settings. In the other settings, *StabCell* showed empirical PFER control, especially in the case with only one true cell type, where no true marker genes are present (Supplementary Fig. S12). Depending on the PFER_max_ threshold, double dipping and Countsplit produced more or similar numbers of false positive markers. TN Test generated the most false positives by far. ClusterDE generally yielded low numbers of false positives. Double dipping, ClusterDE and TN Test were able to recover more or similar numbers of true positives compared to *StabCell* in several settings (Supplementary Fig. S14).

In 10 cell type scenarios with little information, all methods have low ARI values (Supplementary Fig. S17). In these settings, *StabCell* also yields low cell stability scores (Supplementary Fig. S16) and is able to reduce the number of clusters by merging some spurious clusters (Supplementary Fig. S15). If sufficient information is available to differentiate between cell types, *StabCell*, double dipping, and ClusterDE (which uses the same clustering as double dipping) perform similarly in terms of ARI. Countsplit and TN Test show worse clustering performance since they can only use half of the available information in terms of counts or cells, respectively.

If only one true cell population is present, other approaches consistently find more spurious clusters for all tested library sizes (double dipping and ClusterDE: between 7 and 11; Countsplit: between 7 and 12; TN Test: between 6 and 10), whereas *StabCell* consistently detects fewer clusters (between 2 and 9) and therefore reduces overclustering, while failing to reduce the clustering to a single cluster (Supplementary Fig. S15). Observed mean cell stability scores are low in these scenarios (Supplementary Fig. S16). Note that these results depend on the clustering function *f*_*c*_ and the initial clustering *η*^(0)^. Our results illustrate that *StabCell* can add robustness to potentially misspecified analysis pipelines.

### 3.2 Human Cardiomyocyte Differentiation

To evaluate *StabCell*’s performance on real-world data, we analyzed a scRNA-seq dataset of IPSCs differentiating into cardiomyocytes collected over a differentiation time course of 16 days (Elorbany et al., 2022). Two distinct trajectories have been identified. scRNA-seq data from 19 human cell lines were sampled at seven unique time points using Drop-seq (Supplementary Fig. S18). Preprocessed count data are available via the GEO database (accession number GSE175634) and consist of 230,786 cells and 38,943 genes. The same dataset has previously been analyzed by Neufeld et al. (2024b) using Countsplit to identify genes correlated with the trajectories. Therefore, it serves as a natural dataset for comparison, even though our analysis focuses on differential expression analysis rather than trajectory analysis. Since ClusterDE and TN Test are designed for testing a single cell subpopulation against another single subpopulation whereas we are interested in testing one subpopulation against all other subpopulations, we only compare *Stab-Cell* to double dipping and Countsplit on this dataset.

To generate an initial clustering *η*^(0)^, we used a pipeline similar to Elorbany et al. (2022) (implemented entirely in R) with a resolution parameter in the Louvain algorithm that resulted in additional cell subpopulations (Fig. 5a-c). Using the annotations from Elorbany et al. (2022) as a reference, we considered these additional clusters as potentially spurious, resulting from overclustering. Starting with the double dipping clustering as *η*^(0)^, we applied *StabCell* using the same clustering procedure and the Wilcoxon testing procedure in *B* = 50 complementary pairs of subsamples (100 subsamples in total), assumed *r*-concavity, and set PFER_max_ = 100 per cluster.

**Figure 5.**
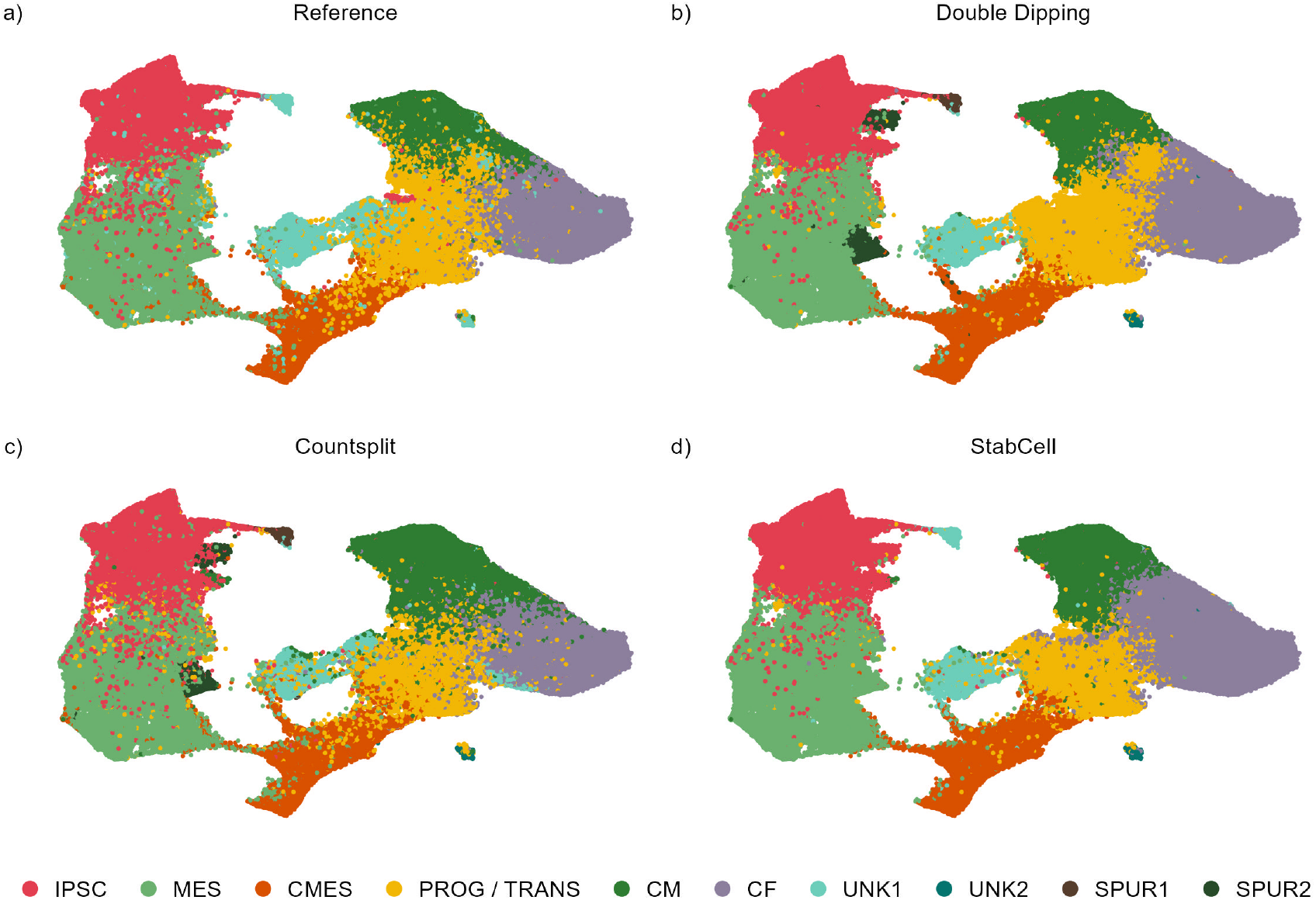
UMAP plots (McInnes et al., 2020) of the cardiomyocyte differentiation dataset. a) Clusters as published by Elorbany et al. (2022). b) Clusters as determined by the double dipping approach with additional, potentially spurious clusters. c) Clusters as determined by Countsplit with additional, potentially spurious clusters. d) Stable clusters as determined by *StabCell*. Cluster labels are induced pluripotent stem cell (IPSC), mesoderm (MES), cardiac mesoderm (CMES), progenitor cells or transition cluster (PROG / TRANS), cardiomyocyte (CM), cardiac fibroblast (CF), unknown (UNK*), and additional, potentially spurious clusters (SPUR*).

Although the initial clustering contained additional, potentially spurious clusters, we observed that these clusters were merged into other clusters (Fig. 5d). The cluster “SPUR1” was merged into cluster “UNK1”, whereas cells from “SPUR2” were reassigned to clusters “IPSC” and “MES”. These changes are in agreement with the results by Elorbany et al. (2022). Furthermore, we observed that cells that were assigned to cluster “PROG / TRANS” had only moderate stability scores, while the other cells were assigned to a single cluster more consistently (Supplementary Fig. S19). Inspecting the individual subsample clusterings showed that the respective cells were sometimes assigned to clusters “CM” and “CF” (Supplementary Fig. S20, e.g., subsample 2).

The number of selected marker genes per cluster (Supplementary Tab. S1) reveals that *StabCell* consistently selected fewer genes than the double dipping approach and Countsplit, which is the result of aggregating the subsample marker sets and only reporting the genes that have been selected most frequently. Inspecting these selection frequencies (Supplementary Fig. S21) shows that most genes either had a very high or a very low selection frequency across all clusters, with at most 16.4% of genes having selection frequencies between 10% and 90%. Some clusters (e.g., clusters “PROG / TRANS” and “CM”) showed patterns of genes that accumulated at certain selection frequencies other than the extremes. This likely reflects inconsistent assignments of cells to different clusters, which was already apparent in the stability scores.

Selected known lineage-associated marker genes are used to assess cluster annotations (Supplementary Fig. S22). Our results agreed with most of the cell type assignments of Elorbany et al. (2022), but we found only limited evidence for the proposed progenitor cells, with the gene ISL1 ranked at position 23–37 across all methods. Conversely, we observed weakly up-regulated expression of genes VIM, RGS5, MDK, and PDGFRA suggesting mesenchymal or transition activity. This is in line with the observation that this cluster was generally less stable and its cells did not show high cell assignment frequencies. Therefore, we think a clear assignment of progenitors would (although biologically plausible) imply unwarranted confidence. Hence, we assigned the mixed cluster label “PROG / TRANS”. For clusters “UNK1” and “UNK2” we again follow Elorbany et al. (2022) and assign an “unknown” label. Our analysis suggests endodermal-like epithelial characteristics for “UNK1” and vascular endothelial-like characteristics for “UNK2”. These clusters may represent alternative differentiation outcomes (Selewa et al., 2020).

To determine the top marker genes for each cluster by method, we considered only upregulated genes, i.e., genes with positive log-fold change (LFC), and ranked them by decreasing selection frequency for *StabCell* and increasing *p*-value for double dipping and Countsplit. Potential ties were ranked by decreasing LFC. While *StabCell* itself does not provide *p*-values, the testing method used on the subsamples does. Genes with repeatedly low *p*-values have a high selection frequency. The selection frequencies can therefore reduce the influence of individual subsample *p*-values on the final marker selection. Comparing the ranks of the marker genes between the different approaches indeed showed that *StabCell* ranked several selected biologically meaningful genes as top markers, while the same genes tended to have higher rank numbers when using double dipping or Countsplit (Tab. 1). The ranking differences may partly be attributable to the different criteria used and to different cluster assignments between the methods. A comparison between the top markers per cluster and method is available in Supplementary Tab. S2.

## 4 Discussion

*StabCell* mitigates the effects of double dipping in scRNA-seq analyses by integrating clustering and differential expression testing within a subsampling framework. Unlike other approaches, it provides cell-specific stability scores and stable marker sets, while allowing arbitrary methods for clustering and marker detection.

**Table 1.** Ranks of selected biologically meaningful marker genes for each cluster (lower values indicate higher-ranked genes). *StabCell* ranks biologically meaningful genes among the top-ranked genes for most clusters. The same genes generally have higher rank numbers when using other approaches.

| Cell Type (Cluster Label) | Marker Gene | *StabCell* | Double Dipping | Countsplit |
|---|---|---|---|---|
| IPSC (IPSC) | L1TD1 | 1 | 72 | 96 |
| mesoderm (MES) | MIXL1 | 6 | 36 | 32 |
| cardiac mesoderm (CMES) | CRABP2 | 1 | 94 | 96 |
| cardiac mesoderm (CMES) | HAS2 | 2 | 63 | 67 |
| cardiac mesoderm (CMES) | SERPINE2 | 4 | 97 | 89 |
| progenitor / transition (PROG / TRANS) | ISL1 | 37 | 23 | 35 |
| cardiomyocyte (CM) | MYL7 | 2 | 50 | 57 |
| cardiac fibroblast (CF) | COL3A1 | 1 | 30 | 52 |
| endodermal/epithelial-like (UNK1) | TTR | 5 | 20 | 12 |
| endothelial-like (UNK2) | KDR | 4 | 80 | 77 |
| endothelial-like (UNK2) | EGFL7 | 9 | 81 | 78 |

We evaluated *StabCell* in simulation studies, showing that the observed mean numbers of false positive markers remained below the specified PFER thresholds in all investigated settings for the “none” and “unimodal” assumptions and in most settings with the “*r*-concavity” assumption. Cell stability scores additionally provide information about the robustness of cluster assignments. In a real data analysis of cardiomyocyte differentiation, we were able to show that potentially spurious clusters can be merged and that meaningful markers are ranked higher with *StabCell* compared to other approaches, which highlights *StabCell*’s robustness and improved interpretability.

One limitation of our method is the increased computational complexity resulting from clustering and marker detection in many subsamples. With modern hardware, parallelization can substantially reduce wall-clock time. In fact, the algorithm is embarrassingly parallel, which allowed us to analyze a large scRNA-seq dataset with more than 230,000 cells and 38,000 genes (Elorbany et al., 2022). With parallelization, *StabCell* showed competitive runtimes compared to alternative methods in a small benchmark study. Furthermore, the reference clustering of the full dataset plays a crucial role. By comparing the stable clustering with the initial reference clustering, we showed that *StabCell* can merge potentially spurious clusters to some extent.

Other methods addressing the double dipping problem are, e.g., Countsplit (Neufeld et al., 2024b), ClusterDE (Song et al., 2026), and TN Test (Zhang et al., 2019). These methods primarily focus on post-clustering inference. *StabCell*, however, combines stable clustering and marker gene selection while additionally providing cell-specific stability scores. Moreover, several competing methods make distributional assumptions about the gene expression data. In contrast, the *StabCell* framework does not itself impose a specific distributional model and can, in principle, be combined with arbitrary clustering and marker detection methods. Furthermore, ClusterDE and TN Test are intended to compare one cluster with one other cluster, whereas *StabCell* offers the flexibility of other testing setups like comparing one cluster with all other clusters. Finally, the other methods aim to control the family-wise error rate (FWER) (with Bonferroni adjustment) or false discovery rate (FDR). *StabCell*, however, aims to control the per-family error rate (PFER).

Our real-data evaluation is limited to one large-scale differentiation dataset, and stability under cell subsampling does not necessarily imply reproducibility across independent biological samples. We have not yet investigated the influence of different quality control (QC) steps in the analysis pipeline, since the datasets from Elorbany et al. (2022) and our simulations were already cleaned or did not require cleaning. Steps such as filtering out low-quality cells (e.g., due to lysis or multiplets) can substantially affect the analysis and the results (Luecken and Theis, 2019). We hypothesize that *StabCell* may also be more robust against QC misspecification, but this remains to be investigated. We also plan to investigate *StabCell*’s performance when integrating multiple datasets to obtain stable markers between, e.g., different experimental conditions.

Overall, *StabCell* can support more robust and reproducible scRNA-seq analyses by providing subsample-based stable clusterings and marker genes, with empirical per-family error rate (PFER) generally remaining below the targeted levels in simulations and biologically meaningful markers prioritized in real data.

## Code and Data Availability

An implementation of *StabCell* in the statistical programming language R is available at https://github.com/LuckyLueck/StabCell. Code to reproduce the results is available at https://github.com/LuckyLueck/StabCell_paper. The analyzed cardiomyocyte differentiation dataset was first published by Elorbany et al. (2022) and can be downloaded from the GEO database with accession number GSE175634.

**Key Points**

- Double use of data for clustering and marker detection in scRNA-seq analyses poses a selective inference problem.
- We propose *StabCell*, a stability-selection-based framework for differential expression analysis, which repeatedly analyzes random subsamples.
- In addition to selecting stable marker genes, *StabCell* can refine cluster assignments and provide cell-specific stability scores.
- In simulations, *StabCell* generally reduces false positive marker selection compared with conventional double dipping and can be flexibly combined with different clustering and marker detection methods.
- On an IPSC dataset, it reliably selected biologically meaningful genes among the top markers, improving the interpretability of results.

