## Supplemental Material for "StabCell: Stability selection for clustering and marker detection in single-cell RNA sequencing"

#### A. Error Control

Meinshausen and Bühlmann (2010) were the first to introduce the concept of stability selection. Their aim was to identify the set of signal variables $S$ and to separate them from the noise variables $N$, where $\hat{S}$ is an estimator of $S$. By subsampling a $d$-dimensional dataset with $n$ observations and further applying any base variable selection procedure to the subsamples, they calculated variable selection frequencies and only selected variables with high selection frequency as stable variables. Assuming that the base selection procedure is better than random sampling and exchangeability of the noise-variable selection indicators yields

$$\text{PFER} = \mathbb{E}\left[\left|\hat{S}^{(\text{stable})} \cap N\right|\right] \leq \frac{1}{2\tau - 1}\frac{q^2}{d}$$

as an upper bound for the per-family error rate (PFER) where $q$ is the expected number of selected variables in a subsample and $\tau \in (0.5, 1)$ is the selection frequency threshold. This framework provides error control for variable selection even if the base selection procedure does not.

Shah and Samworth (2013) extended the theory without relying on a fixed distinction between signal and noise variables. Based on a threshold $\theta \in [0, 1]$ they divide all variables into sets of high selection probability $H_\theta = \{j : \pi_{j,\lfloor \frac{n}{2} \rfloor} > \theta\}$ and low selection probability $L_\theta = \{j : \pi_{j,\lfloor \frac{n}{2} \rfloor} \leq \theta\}$ where $\pi_{j,\lfloor \frac{n}{2} \rfloor}$ is the selection probability of variable $j$ in a random subsample of size $\lfloor \frac{n}{2} \rfloor$ using the base selection procedure. Considering complementary pairs of such subsamples and the probability of a variable to be selected in both of the complementary subsets, they were able to derive three bounds for the number of selected variables with low selection probability $\mathbb{E}\left[\left|\hat{S}^{(\text{stable})} \cap L_\theta\right|\right]$. Hofner et al. (2015) argue that controlling the *expected number of selected variables with low selection probability* can be interpreted similarly to controlling the PFER. Note that genes with low selection probabilities do not necessarily coincide with truly non-differentially expressed genes. The different bounds derived by Shah and Samworth (2013) are:

1. Without making any additional assumption, they obtained the bound

$$\mathbb{E}\left[\left|\hat{S}^{(\text{stable})} \cap L_\theta\right|\right] \leq \frac{\theta}{2\tau - 1}\mathbb{E}\left[\left|\hat{S}^{(\text{b})} \cap L_\theta\right|\right]$$

for $\tau \in (0.5, 1]$, which yields the same numerical upper bound as Meinshausen and Bühlmann (2010) derived for non-complementary subsamples if $\theta = q/d$ and $q = \mathbb{E}\left[\left|\hat{S}^{(b)}\right|\right]$.

2. A tighter bound is available under the assumption that the simultaneous selection frequencies, i.e., the proportion of complementary pairs in which a variable is selected in both subsets, for all $j \in L_\theta$ have a unimodal distribution and if additionally $\theta \leq 1/\sqrt{3}$ holds:

$$\mathbb{E}\left[\left|\hat{S}^{(\text{stable})} \cap L_\theta\right|\right] \leq C(\tau, B)\theta q,$$

with

$$C(\tau, B) = \begin{cases} \frac{1}{4\tau - 2 - 1/B} & \text{if } \tau \in \left(\min\left\{\frac{1}{2} + \theta^2, \frac{1}{2} + \frac{1}{2B} + \frac{3}{4}\theta^2\right\}, \frac{3}{4}\right] \\ \frac{4 - 4\tau + 2/B}{1 + 1/B} & \text{if } \tau \in \left(\frac{3}{4}, 1\right] \end{cases}.$$

3. The tightest bound can be obtained under an $r$-concavity assumption for the selection frequencies ($r = -1/4$) and for the simultaneous selection frequencies ($r = -1/2$) for all $j \in L_\theta$:

$$\mathbb{E}\left[\left|\hat{S}^{(\text{stable})} \cap L_\theta\right|\right] \leq d \min\left\{D(\theta^2, 2\tau - 1, B, -1/2), D(\theta, \tau, 2B, -1/4)\right\},$$

which is valid for $\tau \in (\theta, 1]$ and where $D(\eta, t, B, r)$ denotes the maximum of $\mathbb{P}(X \geq t)$ over all $r$-concave random variables supported on $\{0, 1/B, 2/B, \ldots, 1\}$ with $\mathbb{E}[X] \leq \eta$.

In *StabCell* we estimate a comparison-specific $q$ by calculating the mean number of selected variables, i.e., the marker genes, across all subsamples. A natural choice for $\theta$ is $q/d$, which defines $L_\theta$ as the set of genes with at most average selection probability. By specifying a nominal $\mathrm{PFER}_{\max}$ target it is then possible to calculate the corresponding selection frequency threshold $\tau$.

In some cases, the calculated selection frequency threshold exceeded 1; we then set it to 1, i.e., we still report genes that were selected in all subsamples as stable markers. In such cases, the chosen bound does not support the desired nominal $\mathrm{PFER}_{\max}$ target; however, our simulations only show limited signs of this and selection in all subsamples is a strong indicator of robust differential expression. Whenever such a truncation of $\tau$ occurs, we also flag the results in the provided software implementation.

#### B. Alternative Approach

To avoid information leakage between the subsamples, we propose an alternative approach, where the cells are divided into two splits. One split is used for reference clustering, and cluster labels are transferred to the cells in the other split by $k$-nearest neighbor classification in the principal component (PC) space. The transferred cluster labels can then be used to perform subsample-based marker detection with respect to the independently constructed reference clustering rather than the stable clustering. This alternative variant avoids this information leakage and is more closely aligned with the assumptions underlying complementary-pairs stability selection (Shah and Samworth, 2013). However, it proves to be more conservative, since cells used to construct the reference clustering do not contribute to marker detection (Supplementary Fig. S2). Considering the results from our simulations, the potential dependence between subsamples does not tend to have a severe effect in practice and we still observe approximate empirical PFER control. The increased power to retrieve true positive markers, together with the empirical error control observed in simulations, prompted us to follow the original *StabCell* algorithm. Furthermore, the formation of an intermediate stable clustering $\eta^{(\mathrm{stable})}$ may overcome over-clustering and, therefore, can weaken the selective inference problem.

#### C. Algorithm

**Algorithm S1** *StabCell*: Stable clustering and marker gene selection for single-cell RNA sequencing data

**Input:**
Data $\mathcal{D}$, Number of complementary data splits $B$, Clustering function $f_c$, Test function $f_t$, Initial clustering $\eta^{(0)}$, $p$-value threshold $\lambda$, Stability selection bound assumption, Nominal $\text{PFER}_{\max}$ target

**Output:**
Stable Clustering $\eta^{(\text{stable})}$, Observation stability scores $s_i$, Sets of stable markers $\hat{S}_{l\lambda}^{(\text{stable})}$ for every cluster $l$ in $\eta^{(\text{stable})}$

*# Subsampling*
**for** $c = 1, ..., B$ **do**
  Generate complementary subsamples $\mathcal{D}^{(2c-1)}$ and $\mathcal{D}^{(2c)}$ of size $\lfloor \frac{n}{2} \rfloor$
**end for**

*# Calculate stable clustering*
**for** every subsample $b = 1, \ldots, 2B$ **do**
  Find subsample clustering $\eta^{(b)} = f_c\left(\mathcal{D}^{(b)}\right)$
  **for** every cluster $k \in \eta^{(b)}$ **do**
    Find closest cluster $k' \in \eta^{(0)}$ in terms of Jaccard similarity:

$$k'^{(b)} = \underset{z \in \{1,\ldots,K_0\}}{\arg\max} \frac{\left|\left(\gamma_z^{(0)} \cap I^{(b)}\right) \cap \gamma_k^{(\text{b})}\right|}{\left|\left(\gamma_z^{(0)} \cap I^{(b)}\right) \cup \gamma_k^{(\text{b})}\right|}$$

  **end for**
  Construct aligned clustering $\eta'^{(b)}$ by replacing $k$ with $k'$ in $\eta^{(b)}$
**end for**

**for** every cell $i = 1, \ldots, n$ **do**
  **for** all clusters $k' \in \eta^{(0)}$ **do**
    Calculate assignment frequencies: $\pi_{ik'} = \frac{\sum_{b=1}^{2B} \mathbb{1}\left(i \in I^{(b)},\, \eta_i'^{(b)} = k'\right)}{\sum_{b=1}^{2B} \mathbb{1}\left(i \in I^{(b)}\right)}$
  **end for**
  Determine stable clustering label: $\eta_i^{(\text{stable})} = \arg\max_{z \in \{1,\ldots,K_0\}} \pi_{iz}$
  Calculate observation stability score: $s_i = \frac{\left(\sum_{k'=1}^{K_0} \pi_{ik'}^2\right) - 1/K_0}{1 - 1/K_0}$
**end for**

*# Find stable marker genes in subsamples*
**for** every cluster $l \in \eta^{(\text{stable})}$ **do**
  **for** every subsample $b = 1, \ldots, 2B$ **do**
    Find aligned cluster $l'^{(b)} \in \eta^{(b)}$ with maximal Jaccard similarity

$$l'^{(b)} = \underset{z \in \{1,\ldots,K_b\}}{\arg\max} \frac{\left|\left(\gamma_l^{(\text{stable})} \cap I^{(b)}\right) \cap \gamma_z^{(\text{b})}\right|}{\left|\left(\gamma_l^{(\text{stable})} \cap I^{(b)}\right) \cup \gamma_z^{(\text{b})}\right|}$$

    Using $f_t$, test all variables $j$ for differences between aligned cluster $l'^{(b)}$ and the other observations of data $\mathcal{D}^{(b)}$ and save the Bonferroni corrected $p$-value $p_{lj}^{(b)}$
    Select subsample markers $\hat{S}_{l\lambda}^{(b)} = \left\{ j : p_{lj}^{(b)} \leq \lambda \right\}$
  **end for**
  **for** every gene $j = 1, \ldots, d$ **do**
    Calculate selection frequency $\hat{\Pi}_{lj}^{\lambda} = \frac{1}{2B} \sum_{b=1}^{2B} \mathbb{1}\left(j \in \hat{S}_{l\lambda}^{(b)}\right)$
  **end for**
  Calculate assumption-specific selection frequency threshold $\tau$ as a function of $q$ and $\text{PFER}_{\max}$ (cf. Section A)
  Find sets of stable markers $\hat{S}_{l\lambda}^{(\text{stable})} = \{j : \hat{\Pi}_{lj}^{\lambda} \geq \tau\}$
**end for**

#### D. Tables

**Table S1:** Number of selected marker genes per cluster in the cardiomyocyte dataset. Additional clusters SPUR1 and SPUR2 from the initial clustering were merged in *StabCell*.

| Cluster | *StabCell* (Subsample Mean) | Double Dipping | Countsplit |
|---|---|---|---|
| IPSC | 10662 (14371) | 12378 | 16865 |
| MES | 9988 (12569) | 13560 | 15168 |
| CMES | 5500 (6773) | 8369 | 9103 |
| PROG / TRANS | 7263 (11028) | 10520 | 13101 |
| CM | 7154 (10956) | 10802 | 14361 |
| CF | 10507 (14186) | 14146 | 16073 |
| UNK1 | 3565 (4801) | 6518 | 6409 |
| UNK2 | 1280 (1257) | 1939 | 2066 |
| SPUR1 | | 3090 | 3291 |
| SPUR2 | | 4490 | 7732 |

**Table S2:** Top 20 markers by cluster and method in the cardiomyocyte dataset. *StabCell* markers are ranked by decreasing selection frequency whereas double dipping and Countsplit markers are ranked by increasing Bonferroni-adjusted $p$-value. Ties are broken by decreasing LFC. Only upregulated genes with positive LFC were considered. Selected biologically meaningful marker genes are highlighted.

| Cluster | Rank | *StabCell* | Double Dipping | Countsplit |
|---|---|---|---|---|
| IPSC | 1 | **L1TD1** | MT1G | AC007991.2 |
| | 2 | MIR302CHG | IDO1 | IDO1 |
| | 3 | TERF1 | NPTX1 | NPTX1 |
| | 4 | **DPPA4** | AC007991.2 | MT1G |
| | 5 | **ESRG** | MT1H | ANKRD34B |
| | 6 | POLR3G | AL591030.1 | AL591030.1 |
| | 7 | CD24 | LINC00599 | TNFSF11 |
| | 8 | TMSB4X | CXCL5 | LINP1 |
| | 9 | SFRP2 | LNCPRESS1 | AL138720.1 |
| | 10 | UCHL1 | TRIML2 | LINC00599 |
| | 11 | FOXD3-AS1 | MT1E | MT1H |
| | 12 | THY1 | MT1F | UCMA |
| | 13 | **TDGF1** | UCMA | TRIML2 |
| | 14 | PARP1 | AL138720.1 | LINC02188 |
| | 15 | TKT | MT1X | LNCPRESS1 |
| | 16 | MGST1 | PRDM14 | CXCL5 |
| | 17 | RPL22L1 | FOXD3-AS1 | DPPA5 |
| | 18 | PSIP1 | TRBC2 | PRDM14 |
| | 19 | PTMA | PF4 | MT1F |
| | 20 | UGP2 | ZDHHC22 | PF4 |

Continued on next page.

**Table S2:** (continued)

| Cluster | Rank | *StabCell* | Double Dipping | Countsplit |
|---|---|---|---|---|
| MES | 1 | TUBB2B | WNT10B | AC009654.1 |
| | 2 | L1TD1 | AC022784.2 | WNT10B |
| | 3 | TUBB2A | CSMD1 | LINC01780 |
| | 4 | LINC01356 | LINC01780 | AC022784.2 |
| | 5 | EPCAM | **TBXT** | **TBXT** |
| | 6 | **MIXL1** | GLIPR1L1 | CDX1 |
| | 7 | CNTNAP2 | AC009654.1 | GLIPR1L1 |
| | 8 | MIR302CHG | CDX1 | CSMD1 |
| | 9 | GAS5 | SPATA16 | SPATA16 |
| | 10 | ATP5PD | IL23A | CALCA |
| | 11 | GSTO1 | AL353747.4 | SP5 |
| | 12 | HSPE1 | SP5 | PCSK1 |
| | 13 | IFITM1 | NODAL | AL353747.4 |
| | 14 | AL353747.4 | PCSK1 | NODAL |
| | 15 | SLIRP | EVX1 | IL23A |
| | 16 | DPPA4 | FOXB1 | AL359915.1 |
| | 17 | HSPD1 | WNT5B | FOXB1 |
| | 18 | GAL | TTC29 | EVX1 |
| | 19 | TMSB4X | GAD1 | AC131094.1 |
| | 20 | PMAIP1 | PLA2G2A | DCLK1 |
| CMES | 1 | **CRABP2** | AC092078.2 | AC092078.2 |
| | 2 | **HAS2** | AC005548.1 | CER1 |
| | 3 | **APLNR** | CER1 | AC005548.1 |
| | 4 | **SERPINE2** | LINC01213 | MRAP |
| | 5 | HAPLN1 | MRAP | LINC02192 |
| | 6 | BNIP3 | LINC02192 | LINC01213 |
| | 7 | NTS | **MESP2** | INHBB |
| | 8 | CER1 | RFPL2 | AC117386.2 |
| | 9 | **LHX1** | AC117386.2 | RFPL2 |
| | 10 | KRT19 | BX276092.7 | **MESP2** |
| | 11 | LDHA | POTEC | MAGEA11 |
| | 12 | CDH11 | INHBB | FAM24B |
| | 13 | PKM | FAM24B | AC016074.2 |
| | 14 | GAS5 | MAGEA11 | **MESP1** |
| | 15 | RPL12 | KCNK17 | POTEC |
| | 16 | RPL13A | TBX6 | KCNK17 |
| | 17 | CNTNAP2 | **MESP1** | TBX6 |
| | 18 | **HAND1** | WNT4 | WNT4 |
| | 19 | MSX1 | **APLNR** | BX276092.7 |
| | 20 | HIST1H1D | LGALS12 | LGALS12 |
| PROG / TRANS | 1 | VIM | SPOCK3 | SPOCK3 |
| | 2 | MALAT1 | SIX1 | SIX1 |
| | 3 | LIX1 | EBF2 | HOXB1 |
| | 4 | TPM1 | FOXC2 | FOXC2 |
| | 5 | RHOBTB3 | SPX | ALX1 |
| | 6 | MDK | FGF10 | SMOC1 |
| | 7 | PRTG | HOXB1 | HOTAIRM1 |
| | 8 | RGS5 | ALX1 | PRDM1 |
| | 9 | GATA6-AS1 | NTRK2 | CFC1 |
| | 10 | CCDC34 | SMOC1 | AC022168.1 |
| | 11 | PTPN13 | AC022168.1 | NTRK2 |
| | 12 | TMEM88 | PRDM1 | FGF10 |
| | 13 | MEIS2 | CFC1 | MT-TV |
| | 14 | **PDGFRA** | RARB | HOXB-AS1 |
| | 15 | NID2 | COL2A1 | LIX1 |
| | 16 | CTSV | NKX3-1 | RARB |
| | 17 | AC020909.2 | NRG1 | NR2F1 |
| | 18 | DUSP6 | FYB2 | **PDGFRA** |
| | 19 | COL2A1 | HOXB-AS1 | COL2A1 |
| | 20 | PKP2 | LIX1 | PCOLCE |

Continued on next page.

**Table S2:** (continued)

| Cluster | Rank | *StabCell* | Double Dipping | Countsplit |
|---|---|---|---|---|
| CF | 1 | **COL3A1** | ANXA8L1 | ANXA8L1 |
| | 2 | **DCN** | **DCN** | OLR1 |
| | 3 | TPM1 | OLR1 | **DCN** |
| | 4 | **SPARC** | MGP | CLIC3 |
| | 5 | **LUM** | HPCAL4 | RGCC |
| | 6 | S100A10 | AC245041.2 | AC245041.2 |
| | 7 | **COL1A2** | LINC01638 | AC090044.1 |
| | 8 | VIM | AC097480.1 | NPNT |
| | 9 | MALAT1 | AQP1 | AQP1 |
| | 10 | EZR | PROK1 | AC245041.1 |
| | 11 | IGFBP5 | AC245041.1 | PROK1 |
| | 12 | KRT8 | RGCC | HPCAL4 |
| | 13 | **COL1A1** | NPNT | PRPH |
| | 14 | FRZB | CLIC3 | LINC01266 |
| | 15 | MDK | ANXA8 | MGP |
| | 16 | SEPTIN7 | TYRP1 | UPK3B |
| | 17 | GPC3 | RSPO1 | ANXA8 |
| | 18 | HAND1 | AKR1B10 | ABI3BP |
| | 19 | CALD1 | HOXC8 | RSPO1 |
| | 20 | H19 | UPK3B | HOXC8 |
| CM | 1 | EMC10 | ABRA | ABRA |
| | 2 | **MYL7** | **MYH6** | **MYH6** |
| | 3 | **TNNT2** | **MYH7** | **MYBPC3** |
| | 4 | TPM1 | CSRP3 | CSRP3 |
| | 5 | **MYH6** | AC020909.2 | AC020909.2 |
| | 6 | **TTN** | **MYBPC3** | ASB2 |
| | 7 | MYL4 | EMC10 | **MYH7** |
| | 8 | **ACTC1** | HSPB3 | HSPB3 |
| | 9 | TNNI1 | NPPA | ACTA1 |
| | 10 | AC020909.2 | PPP1R3A | EMC10 |
| | 11 | ANKRD1 | CKM | MYOM1 |
| | 12 | NEBL | ASB2 | PPP1R3A |
| | 13 | NEXN | ACTA1 | NPPA |
| | 14 | HSPB1 | MYOM1 | CKM |
| | 15 | MALAT1 | MYL3 | SMPX |
| | 16 | VIM | SMPX | SYNPO2L |
| | 17 | FILIP1 | **ACTC1** | XIRP1 |
| | 18 | ACTA2 | LDB3 | **ACTC1** |
| | 19 | MYL9 | COX6A2 | LDB3 |
| | 20 | TNNC1 | XIRP1 | TCAP |
| UNK1 | 1 | **KRT19** | COL19A1 | LINC02413 |
| | 2 | **APOA1** | SCGB1A1 | COL19A1 |
| | 3 | **APOA2** | LINC00348 | ALAS2 |
| | 4 | FN1 | LINC02413 | LINC00348 |
| | 5 | **TTR** | CGA | CGA |
| | 6 | ID1 | AC020656.2 | AC020656.2 |
| | 7 | PCAT14 | IHH | S100A14 |
| | 8 | **EPCAM** | ALAS2 | CST4 |
| | 9 | APOC1 | OR51B5 | LINC01767 |
| | 10 | S100A14 | LINC00261 | SOAT2 |
| | 11 | ANKRD1 | HABP2 | LINC00261 |
| | 12 | FAM184A | S100A14 | **TTR** |
| | 13 | GATA3 | HNF1B | CST1 |
| | 14 | SLC2A3 | FOXA1 | HNF1B |
| | 15 | DLK1 | CST4 | IHH |
| | 16 | TTC3 | RIPPLY3 | HABP2 |
| | 17 | RGS5 | LINC01767 | RIPPLY3 |
| | 18 | IFI16 | SOAT2 | CST2 |
| | 19 | SERPINE2 | CST1 | FOXA1 |
| | 20 | S100A13 | **TTR** | TTC6 |

Continued on next page.

**Table S2:** (continued)

| Cluster | Rank | *StabCell* | Double Dipping | Countsplit |
|---|---|---|---|---|
| UNK2 | 1 | GNG11 | SOST | PIK3CG |
| | 2 | HAPLN1 | SELE | CLEC1B |
| | 3 | TFPI | CLEC1B | CD93 |
| | 4 | **KDR** | PIK3CG | SELE |
| | 5 | SOST | GIMAP4 | GIMAP8 |
| | 6 | TFPI2 | CD93 | SOST |
| | 7 | IFI16 | F2RL2 | GIMAP4 |
| | 8 | CRHBP | AKR1C4 | HID1-AS1 |
| | 9 | **EGFL7** | **PECAM1** | **PECAM1** |
| | 10 | RDX | GIMAP8 | C2orf66 |
| | 11 | RGS5 | **CDH5** | AL713998.1 |
| | 12 | NRP2 | CD34 | CD34 |
| | 13 | RAMP2 | FCN3 | F2RL2 |
| | 14 | F2R | HID1-AS1 | **CDH5** |
| | 15 | VIM | EMCN | PLVAP |
| | 16 | ITM2B | ESAM | GIMAP1 |
| | 17 | LIMCH1 | DIPK2B | FCN3 |
| | 18 | ID3 | GIMAP1 | GPR182 |
| | 19 | ID1 | GPR182 | ESAM |
| | 20 | **FLT1** | AL713998.1 | EMCN |
| SPUR1 | 1 | | AFP | AFP |
| | 2 | | SERPINA7 | TM4SF4 |
| | 3 | | PLG | ITIH2 |
| | 4 | | ITIH1 | SERPINA1 |
| | 5 | | AHSG | KNG1 |
| | 6 | | FGG | C8B |
| | 7 | | ITIH2 | AHSG |
| | 8 | | SERPINA1 | TINAG |
| | 9 | | KNG1 | ANGPTL3 |
| | 10 | | C8B | FGG |
| | 11 | | SERPINA6 | FGB |
| | 12 | | TINAG | ALB |
| | 13 | | TM4SF4 | APOC3 |
| | 14 | | ALB | FGA |
| | 15 | | APOC3 | SLCO2B1 |
| | 16 | | FGB | ITIH1 |
| | 17 | | SLCO2B1 | GSTA2 |
| | 18 | | FGA | LIPC |
| | 19 | | GSTA2 | PLG |
| | 20 | | LIPC | SERPIND1 |
| SPUR2 | 1 | | MTRNR2L8 | ZNF99 |
| | 2 | | PCSK1 | MTRNR2L8 |
| | 3 | | AC016769.5 | ZNF492 |
| | 4 | | DKK4 | PCSK1 |
| | 5 | | GABRB2 | AC016769.5 |
| | 6 | | TBXT | ZNF560 |
| | 7 | | NODAL | DKK4 |
| | 8 | | AP000459.2 | ZNF208 |
| | 9 | | NEFL | PAX8-AS1 |
| | 10 | | MIXL1 | ZNF726 |
| | 11 | | SNHG5 | TBXT |
| | 12 | | ZFP42 | GABRB2 |
| | 13 | | EPCAM | NODAL |
| | 14 | | CACYBP | LRAT |
| | 15 | | C1QBP | AP000459.2 |
| | 16 | | IFITM1 | NEFL |
| | 17 | | POLR3G | MIXL1 |
| | 18 | | UBE2T | GNG4 |
| | 19 | | CRABP1 | POU5F1B |
| | 20 | | JPT1 | ZFP42 |

### E. Figures

#### E.1. Methods

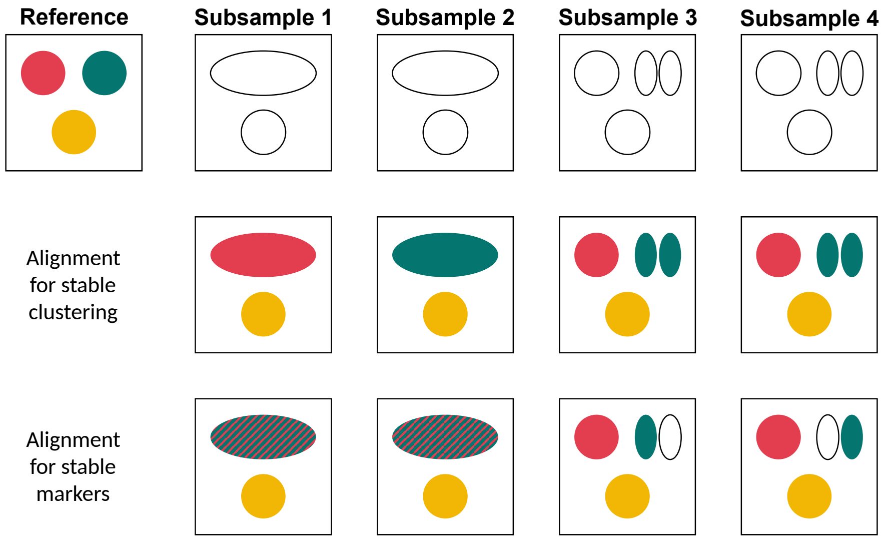


**Figure S1:** Subsample cluster alignment: 1) Alignment for stable clustering: Every cell in the subsample is assigned to a cluster of the reference (here the initial clustering $\eta^{(0)}$). Between subsamples the same cell can be assigned to different reference clusters (Subsample 1 vs 2) or multiple subsample clusters can be aligned to the same reference cluster (Subsample 3 & 4). 2) Alignment for stable markers: For every cluster in the reference clustering (here the stable clustering $\eta^{(\text{stable})}$) an aligned cluster in the subsample is needed to identify corresponding marker genes across subsamples. Multiple reference clusters can be aligned to the same subsample cluster (Subsamples 1 & 2) and some subsample clusters may not be aligned to any reference cluster (Subsamples 3 & 4).

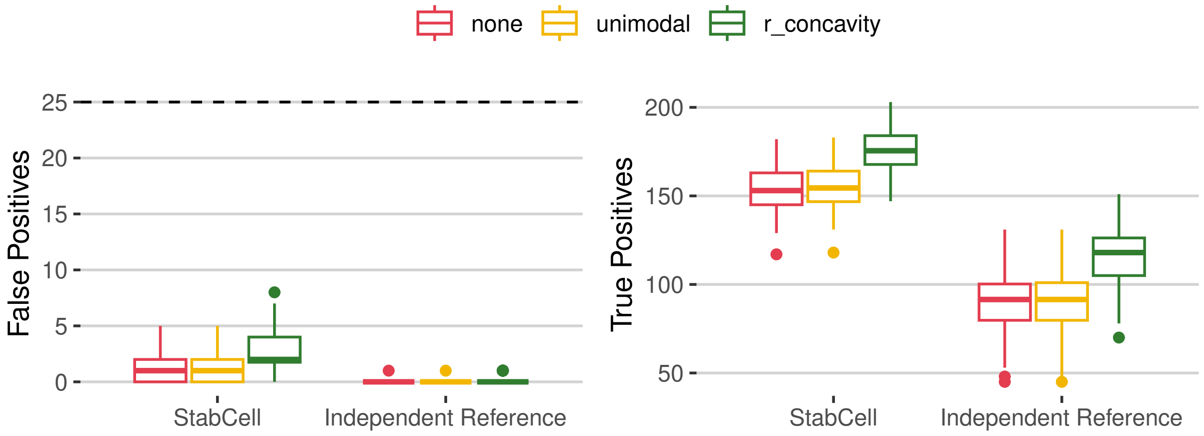


**Figure S2:** We compare the original *StabCell* algorithm ($\text{PFER}_{\text{max}} = 25$) with a variant where the data is split for clustering and marker detection to obtain an independent reference clustering. `R` package `splatter` (Zappia et al., 2017) was used to simulate RNA count data with 5,000 cells and 10,000 genes divided into 10 equally sized subpopulations. Parameters for differential expression are `de.prob` = 0.05, `de.facLoc` = 0.1 and the library size parameter `lib.loc` is set to 11. For the alternative variant the cells are split into sets of equal size. 10-nearest-neighbor classification in the principal component space is used to transfer cluster labels. The shown boxplots are the results of 100 independent simulations and show the numbers of false and true positive markers for the cluster with label 1.

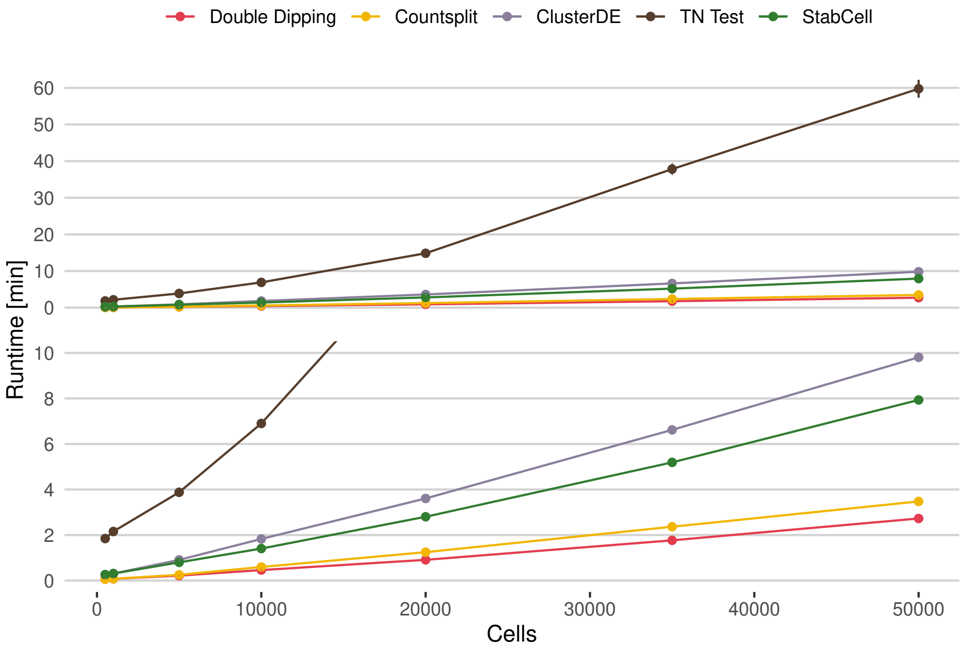


**Figure S3:** Runtime comparison for *StabCell* ($r$-concavity, $\mathrm{PFER}_{\max} = 25$, $B = 50$) and competing methods (top: complete view, bottom: detail view). The code was executed on a system with two AMD EPYC 9654 processors. Parallelization was enabled for *StabCell*, ClusterDE and TN Test with 25 workers and 4 threads. Every simulation had 2 true cell types and 2,500 genes simulated with R package `splatter` (Zappia et al., 2017; `lib.loc`= 9.23, `de.facLoc`= 0.4, `de.prob`= 0.05). Standard `Seurat` functions (Hao et al., 2024) with their default parameters were used for analysis. Shown runtimes are mean values with standard errors determined over 25 independent simulations, but it should be noted that absolute runtimes heavily depend on the chosen clustering and marker detection functions.

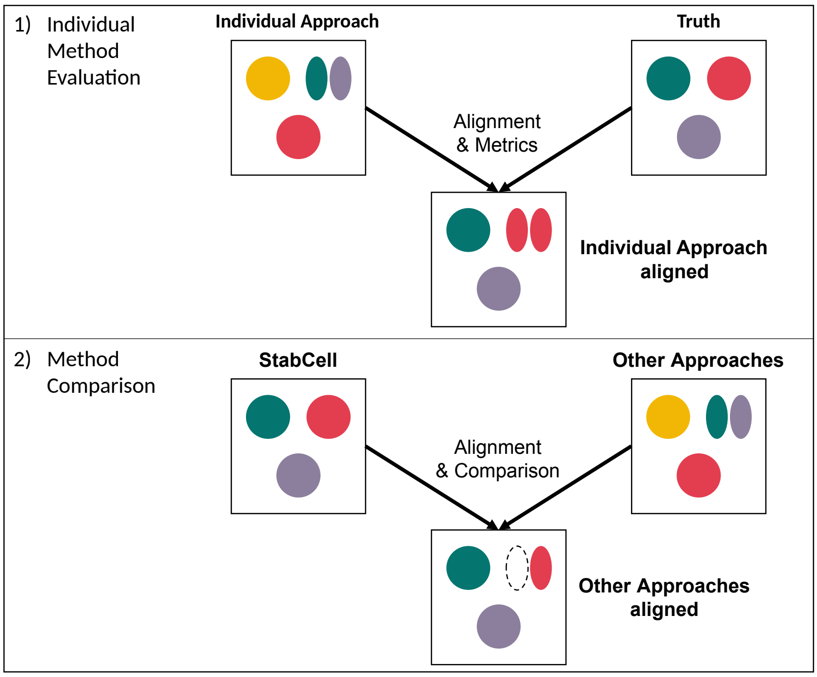


**Figure S4:** Jaccard similarity based cluster alignment for performance evaluation: 1) Each cluster in each approach is aligned to its closest true cluster for metric calculation. Markers between aligned clusters are used to calculate, e.g., false positives. 2) Each *StabCell* cluster and the closest cluster from other approaches is used for method comparisons.

#### E.2. Two Cluster Simulation

*E.2.1. Markers*

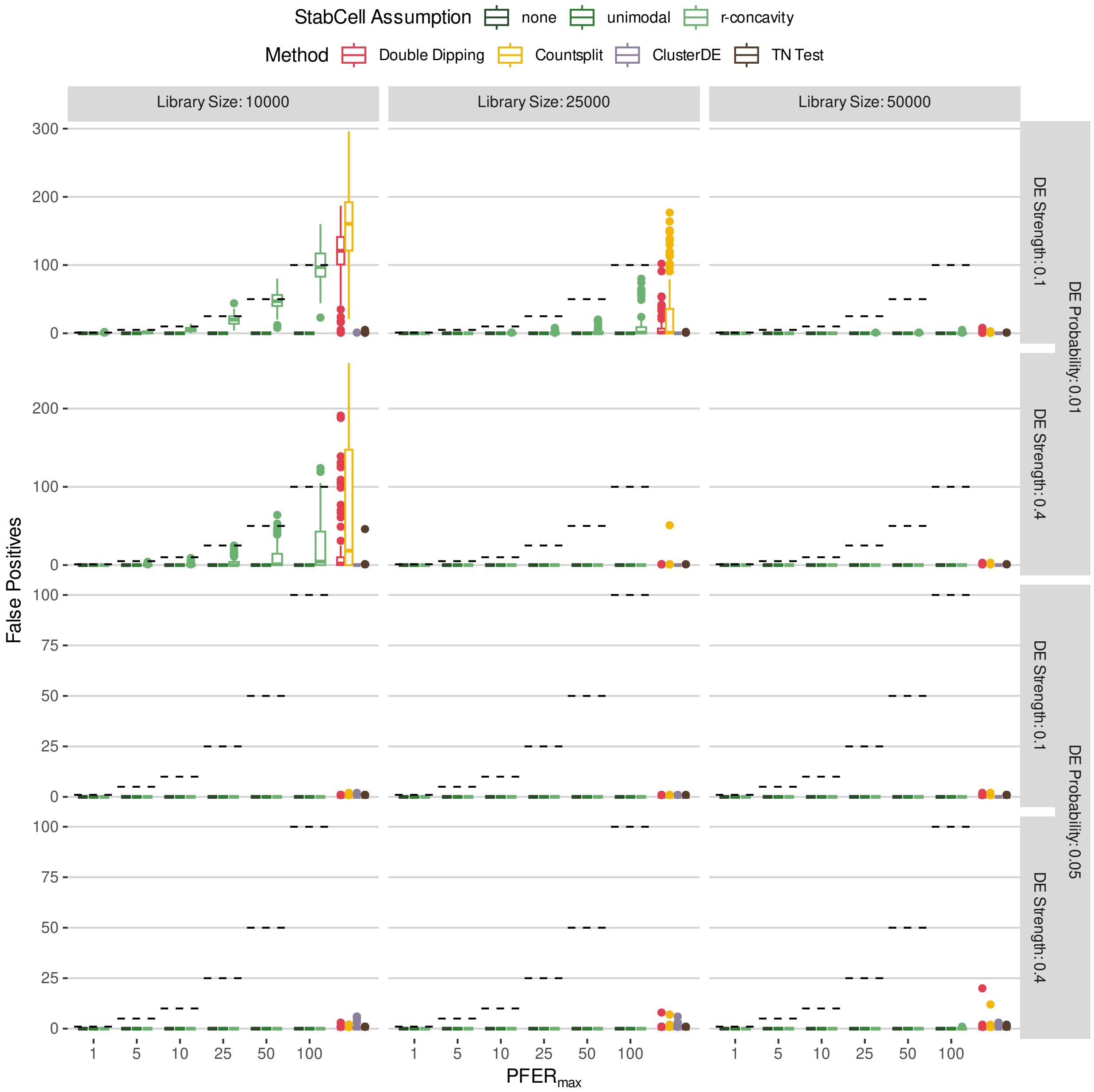


**Figure S5:** False positive results when performing marker detection for 2 true clusters. Boxplots are created using 100 independent repetitions of `splatter` simulations (Zappia et al., 2017; 5,000 cells, 10,000 genes) and *StabCell* analysis with standard Seurat (Hao et al., 2024) functions.

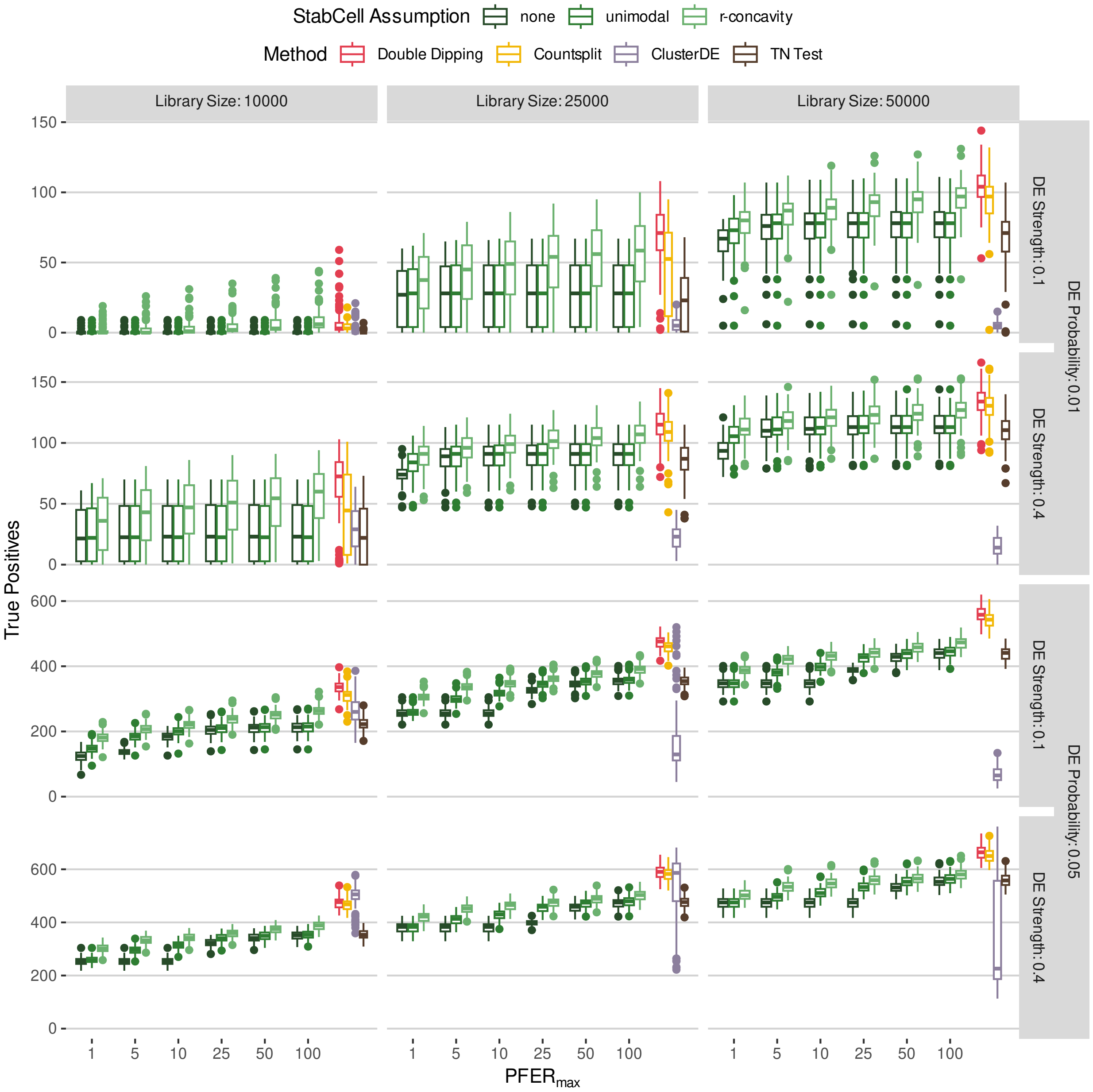


**Figure S6:** True positive results when performing marker detection for 2 true clusters. Boxplots are created using 100 independent repetitions of `splatter` simulations (Zappia et al., 2017; 5,000 cells, 10,000 genes) and *StabCell* analysis with standard Seurat (Hao et al., 2024) functions.

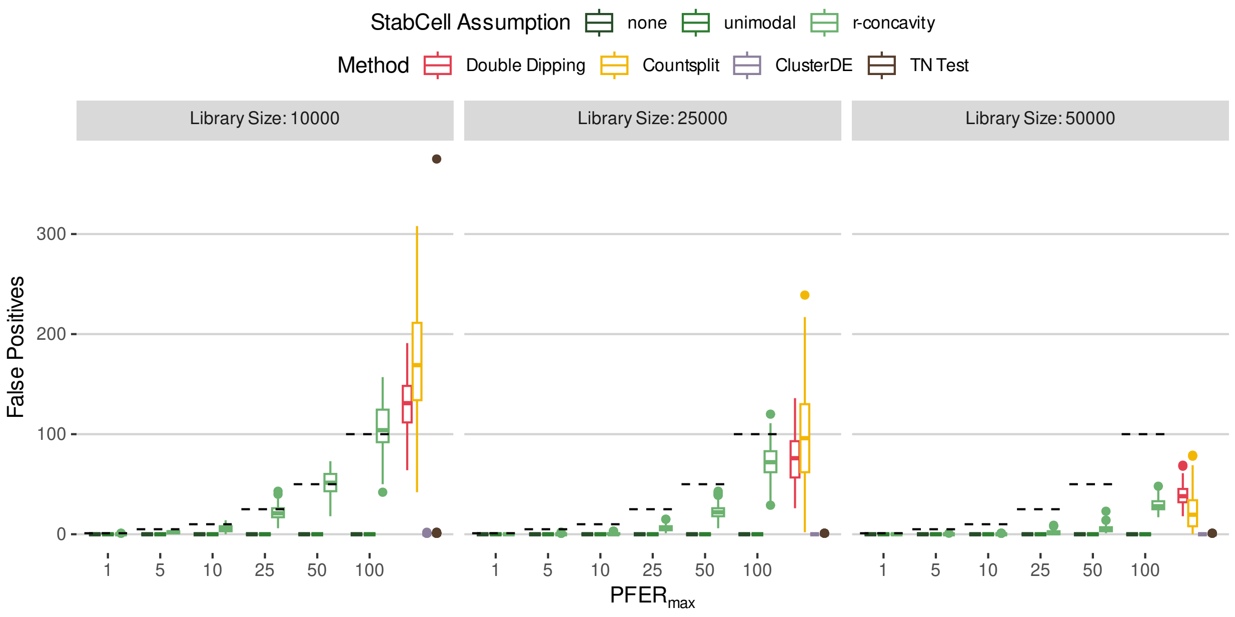


**Figure S7:** False positive results when performing marker detection for 1 true cluster. Boxplots are created using 100 independent repetitions of `splatter` simulations (Zappia et al., 2017; 5,000 cells, 10,000 genes) and *StabCell* analysis with standard Seurat (Hao et al., 2024) functions.

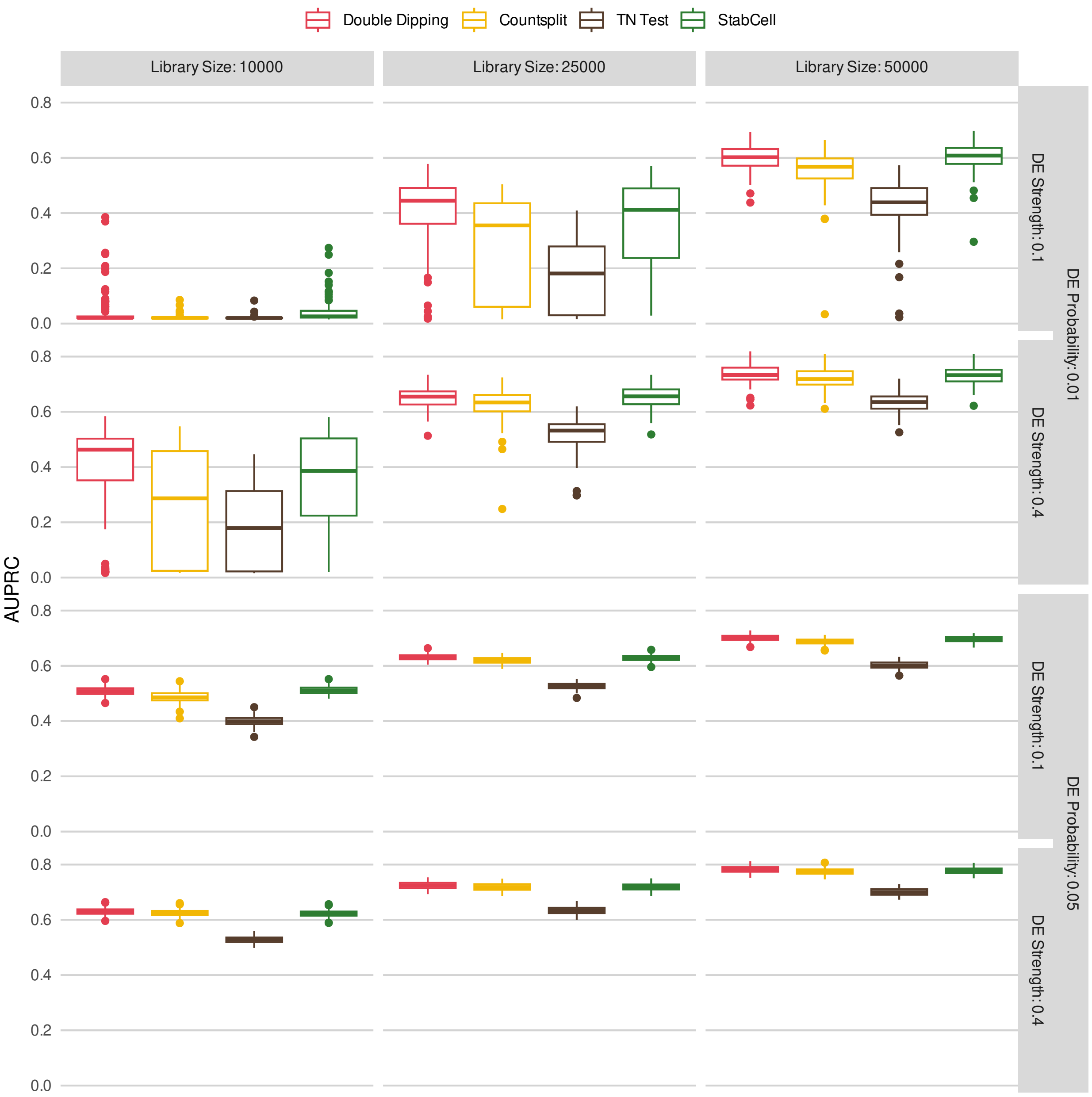


**Figure S8:** Varying *StabCell*'s selection frequency threshold $\tau$ or other methods' $p$-value thresholds yields different sets of selected markers. They are used to calculate the area under the precision-recall curve (AUPRC). Boxplots are created using 100 independent repetitions of `splatter` simulations (Zappia et al., 2017; 5,000 cells, 10,000 genes) and *StabCell* analysis with standard Seurat (Hao et al., 2024) functions.

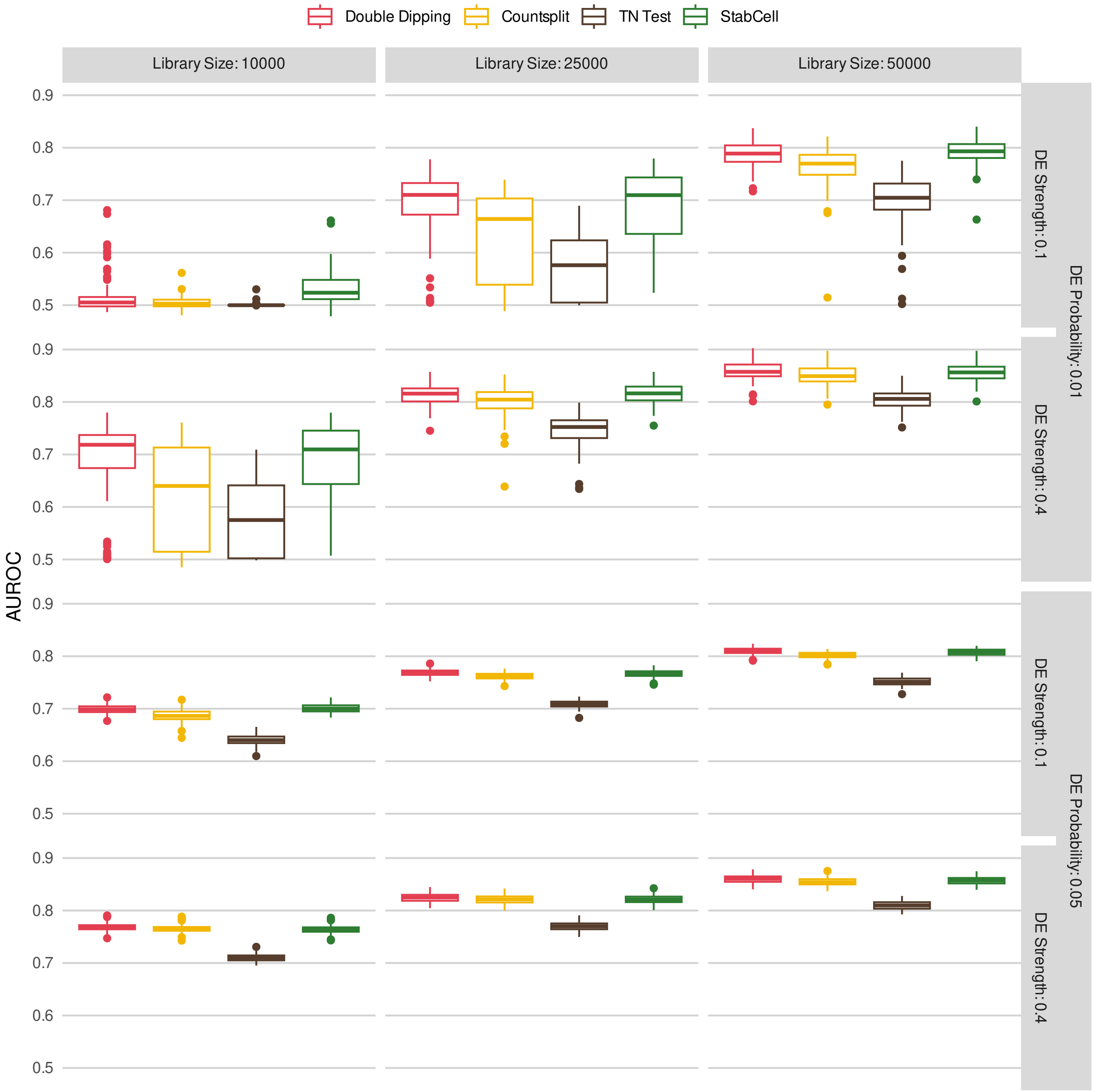


**Figure S9:** Varying *StabCell*'s selection frequency threshold $\tau$ or other methods' $p$-value thresholds yields different sets of selected markers. They are used to calculate the area under the receiver operating characteristic curve (AUROC). Boxplots are created using 100 independent repetitions of `splatter` simulations (Zappia et al., 2017; 5,000 cells, 10,000 genes) and *StabCell* analysis with standard Seurat (Hao et al., 2024) functions.

*E.2.2. Clustering*

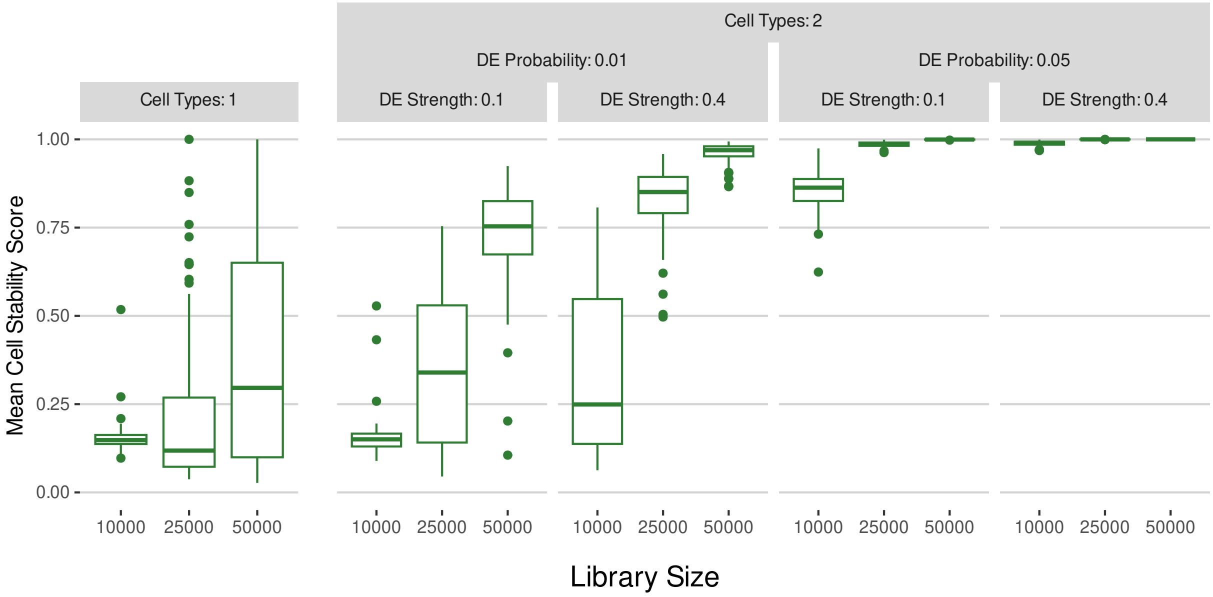


**Figure S10:** Boxplots of the mean cell stability score over all clusters in the 2 cell type scenario. They are created using 100 independent repetitions of `splatter` simulations (Zappia et al., 2017; 5,000 cells, 10,000 genes) and *StabCell* analysis with standard Seurat (Hao et al., 2024) functions.

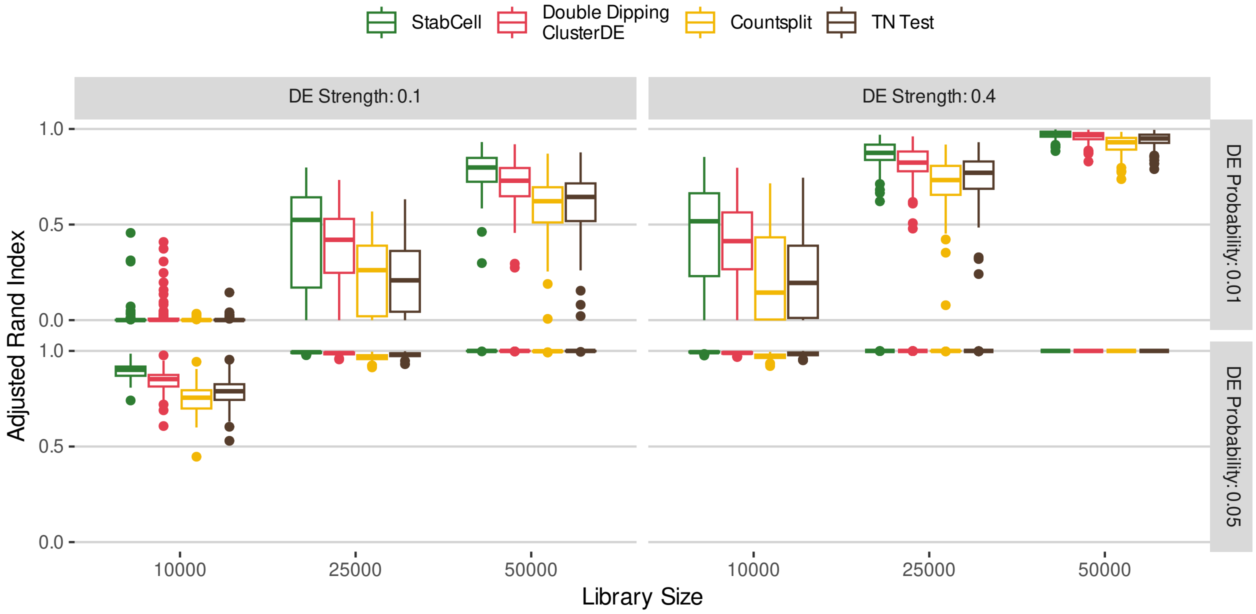


**Figure S11:** Adjusted Rand Index (ARI) results for 2 simulated clusters and all differential expression settings.

#### E.3. Many Cluster Simulation

##### *E.3.1. Markers*

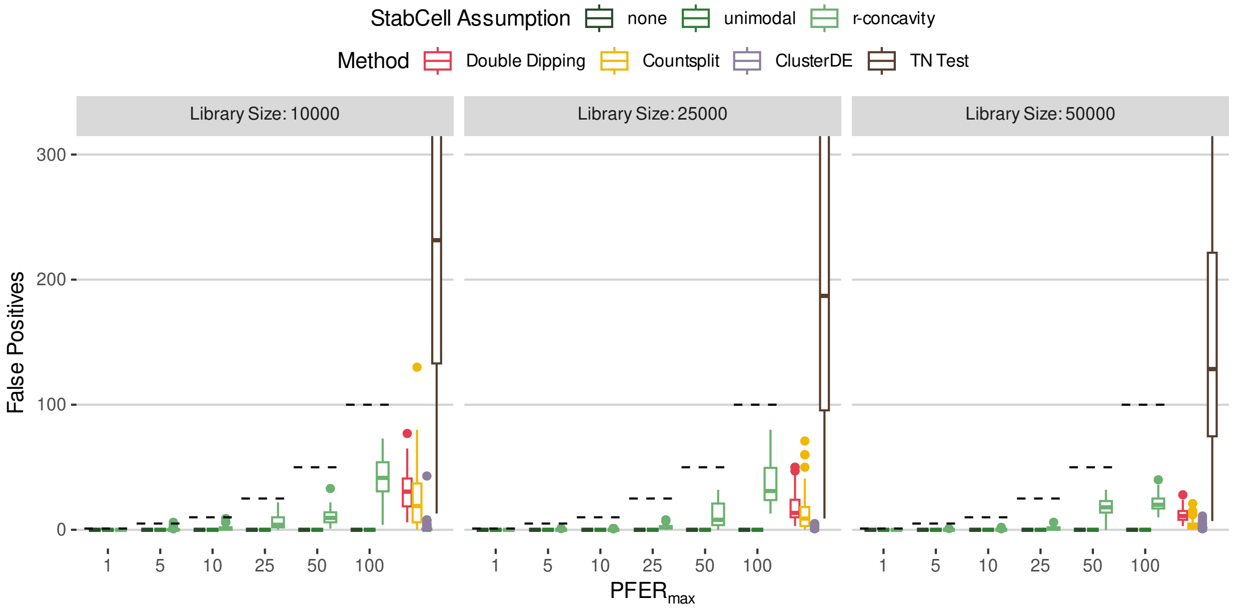


**Figure S12:** False positive results when performing marker detection for 1 true cluster. Boxplots are created using 100 independent repetitions of `splatter` simulations (Zappia et al., 2017; 5,000 cells, 10,000 genes) and *StabCell* analysis with standard Seurat (Hao et al., 2024) functions.

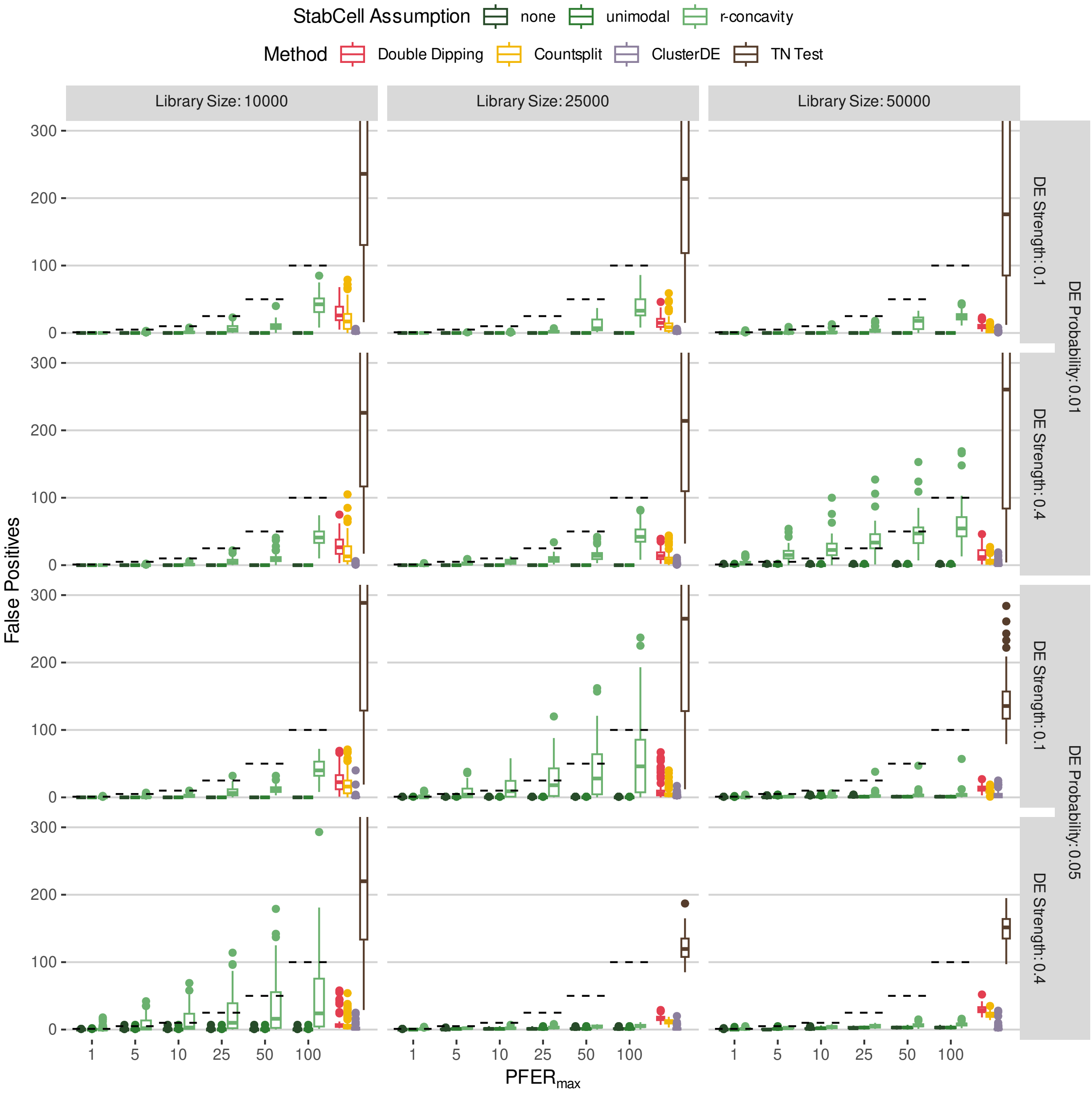


**Figure S13:** False positive results when performing marker detection for 10 true clusters. Boxplots are created using 100 independent repetitions of `splatter` simulations (Zappia et al., 2017; 5,000 cells, 10,000 genes) and *StabCell* analysis with standard Seurat (Hao et al., 2024) functions.

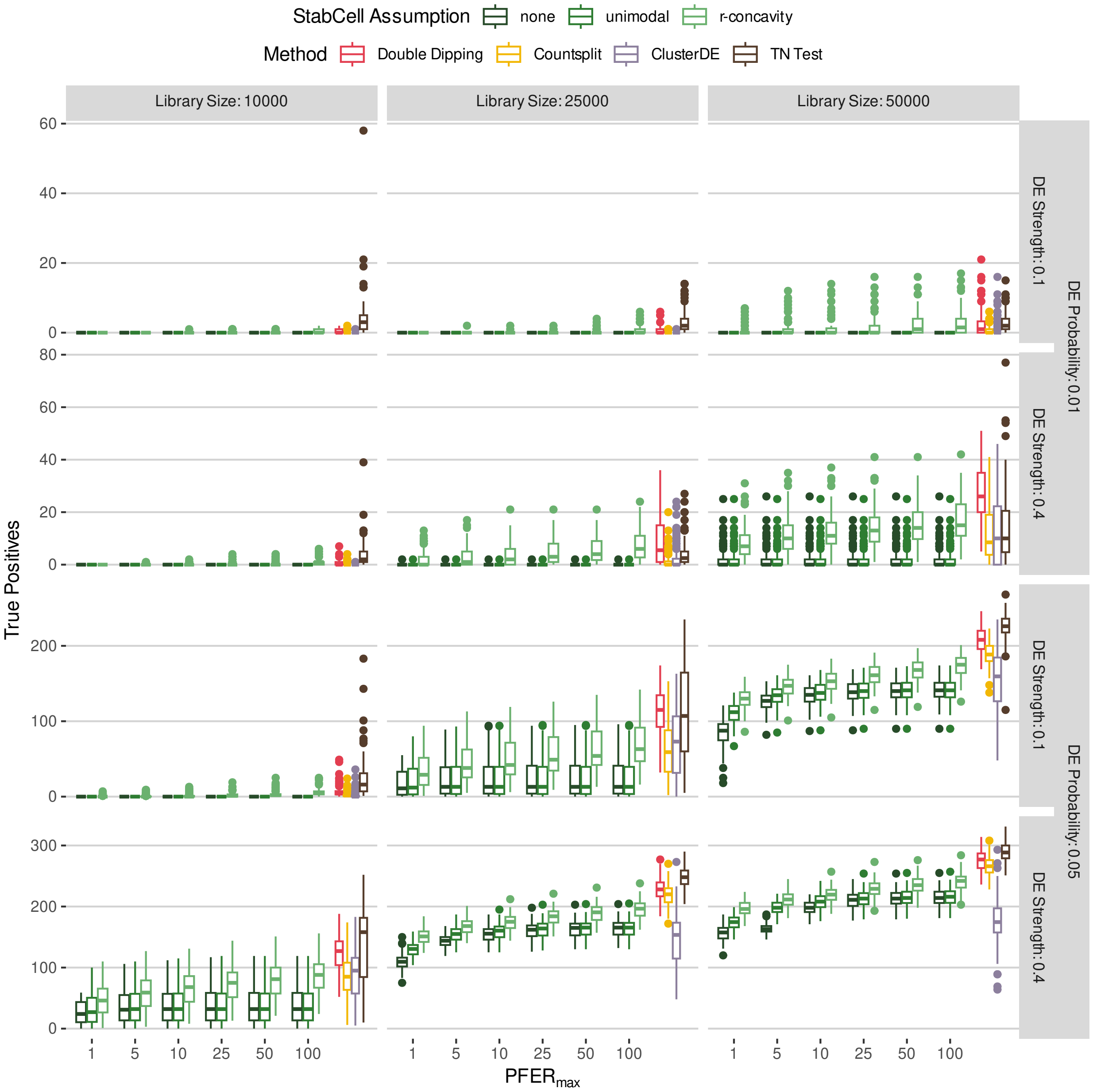


**Figure S14:** True positive results when performing marker detection for 10 true clusters. Boxplots are created using 100 independent repetitions of `splatter` simulations (Zappia et al., 2017; 5,000 cells, 10,000 genes) and *StabCell* analysis with standard Seurat (Hao et al., 2024) functions.

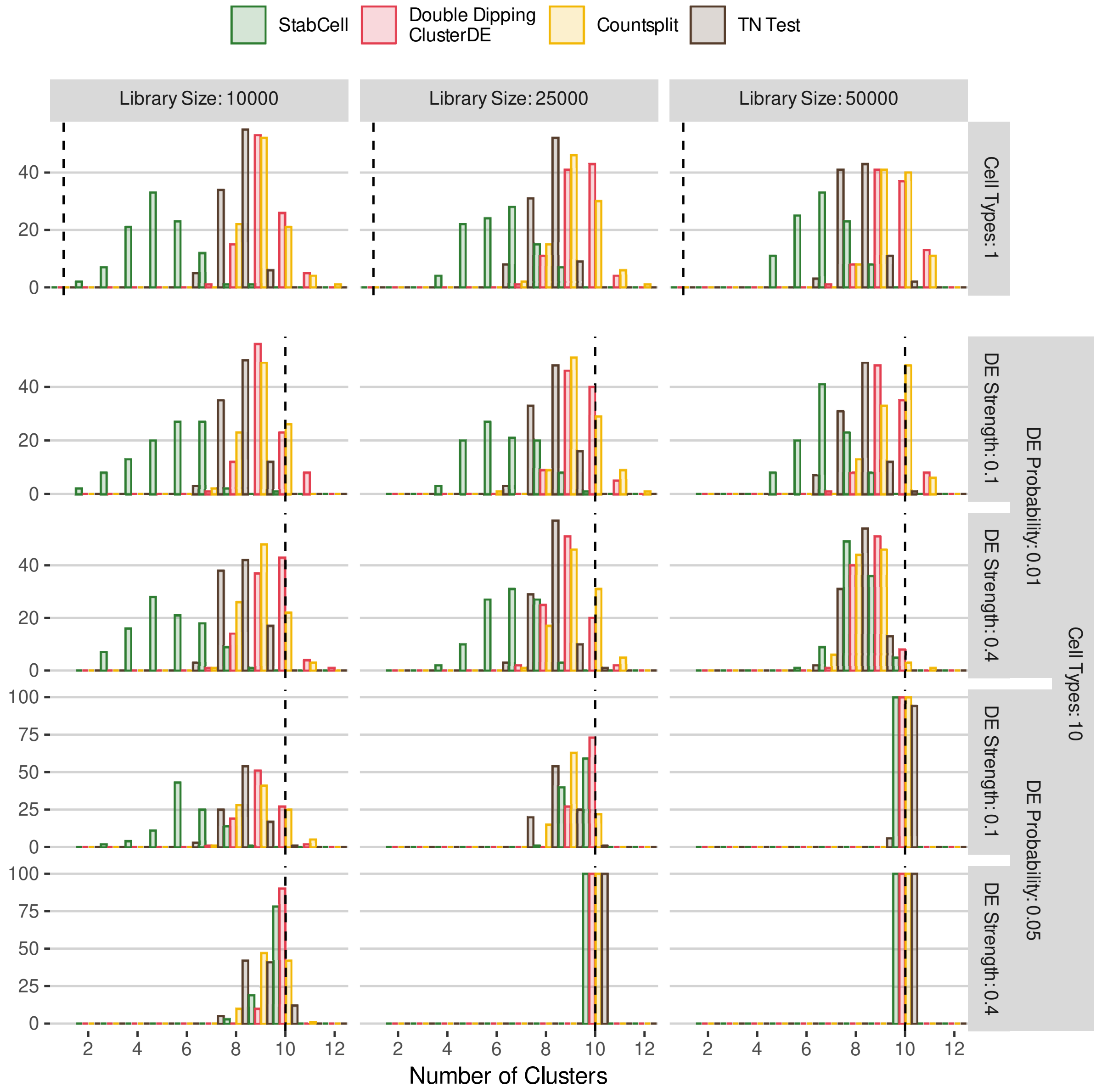


**Figure S15:** Histogram of the obtained number of clusters by method over 100 independent repetitions of `splatter` simulations (Zappia et al., 2017; 5,000 cells, 10,000 genes) and *StabCell* analysis with standard Seurat (Hao et al., 2024) functions. Dashed lines show the true number of simulated cell populations.

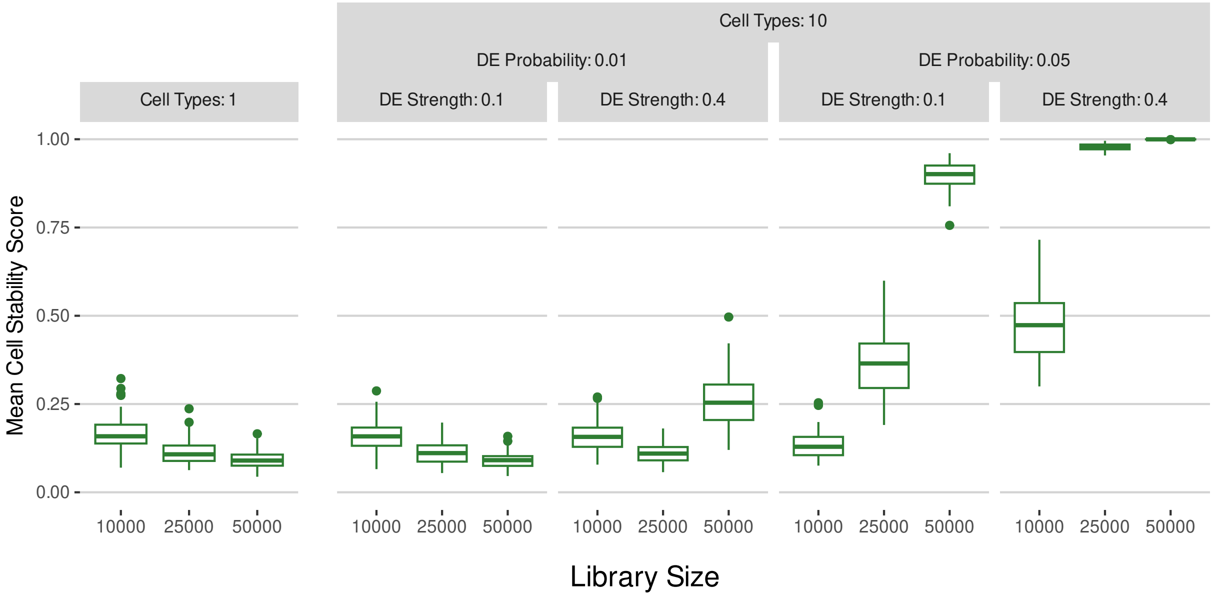


**Figure S16:** Boxplots of the mean cell stability score over all clusters in the 10 cell type scenario. They are created using 100 independent repetitions of `splatter` simulations (Zappia et al., 2017; 5,000 cells, 10,000 genes) and *StabCell* analysis with standard Seurat (Hao et al., 2024) functions.

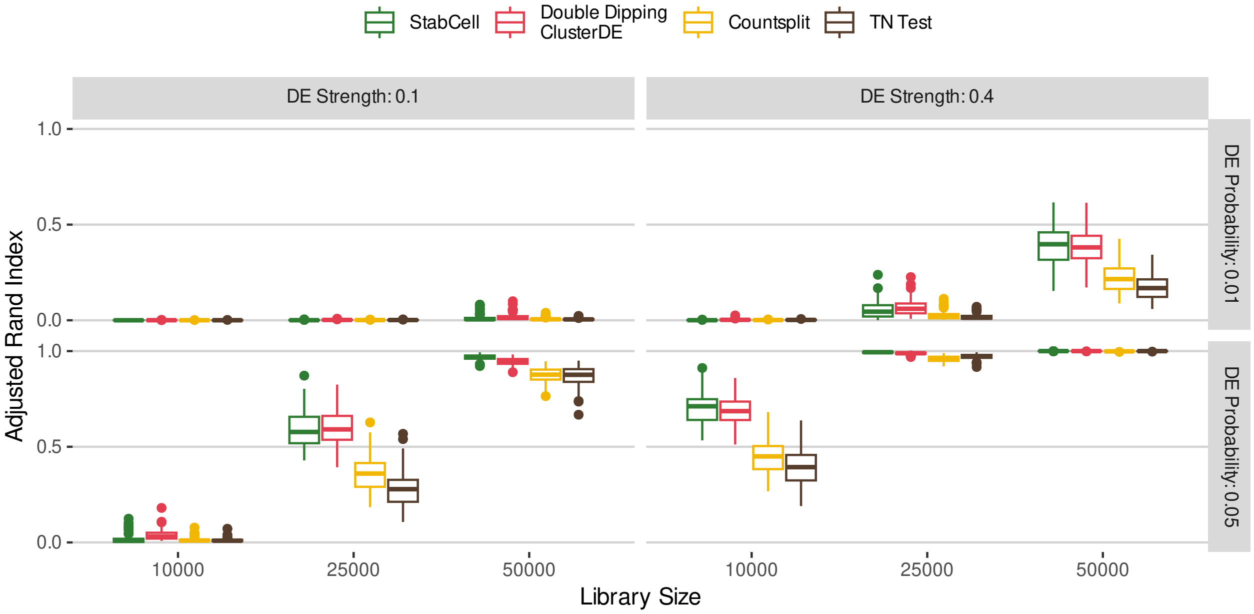


**Figure S17:** Adjusted Rand Index (ARI) results for 10 simulated clusters and all differential expression settings.

E.4. Cardiomyocyte Differentiation

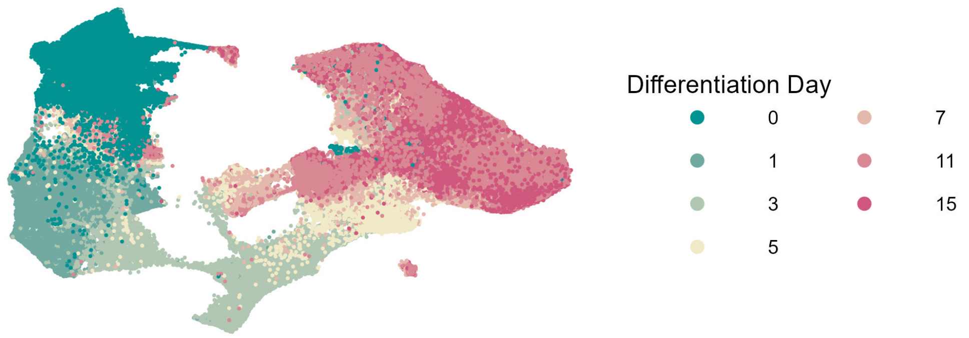


**Figure S18:** UMAP plot (McInnes et al., 2020) of cardiomyocyte dataset colored by differentiation day.

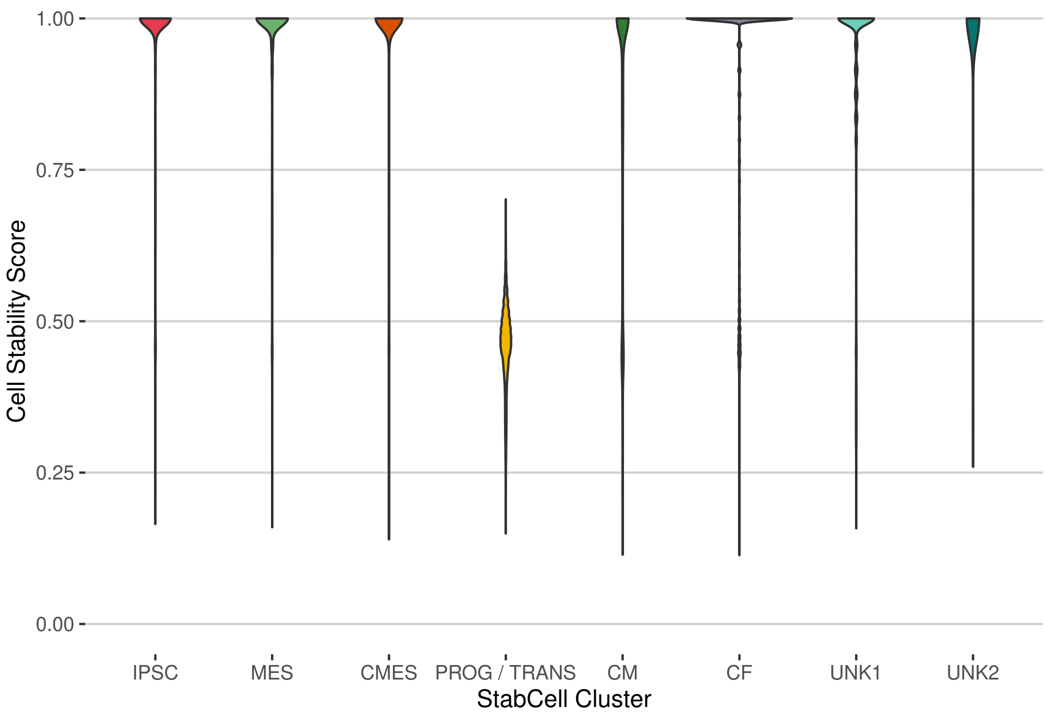


**Figure S19:** Violin plot of cell stability scores per cluster in the cardiomyocyte data.

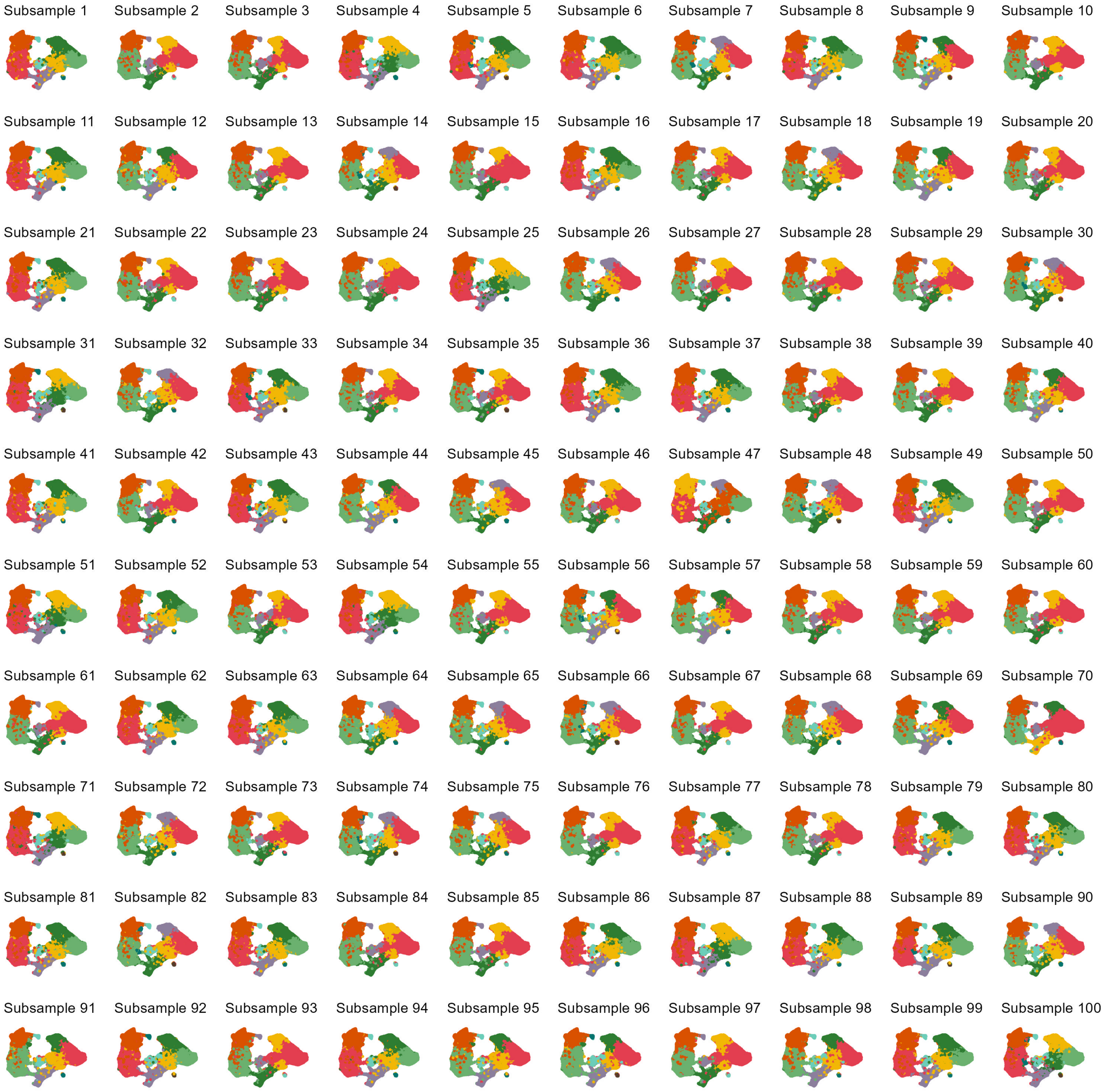


**Figure S20:** Unaligned clusterings of the individual subsamples from the cardiomyocyte dataset.

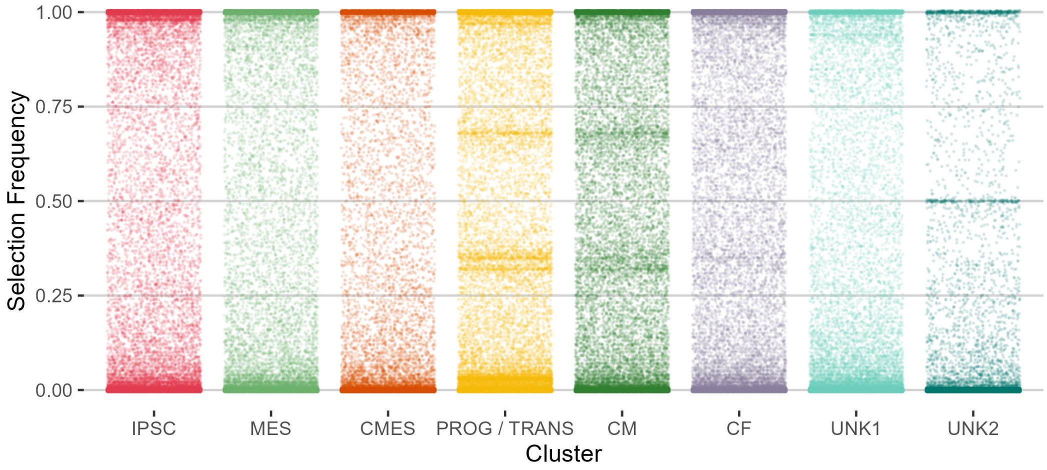


**Figure S21:** Selection frequencies of all genes in all clusters of the cardiomyocyte data. They are most likely to be very high or very low. Every point corresponds to a gene and only genes with a high selection frequency are selected as stable markers.

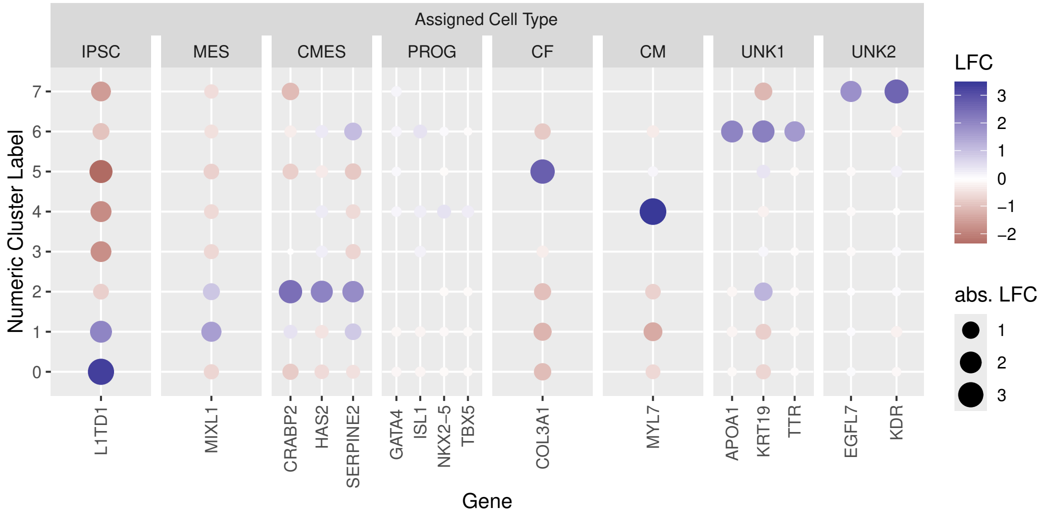


**Figure S22:** Log-fold changes (LFC) of known marker genes support assignment of cell types to clusters in the cardiomyocyte data: 0) induced pluripotent stem cells (IPSC) 1) mesoderm (MES) 2) cardiac mesoderm (CMES) 3) progenitor / transition (PROG / TRANS) 4) cardiomyocyte (CM) 5) cardiac fibroblast (CF) 6) epithelial/endoderm-like (UNK1) 7) endothelial-like (UNK2)
